# A pedigree-based prediction model identifies carriers of deleterious *de novo* mutations in families with Li-Fraumeni syndrome

**DOI:** 10.1101/2020.02.10.942409

**Authors:** Fan Gao, Xuedong Pan, Elissa B. Dodd-Eaton, Carlos Vera Recio, Matthew D. Montierth, Jasmina Bojadzieva, Phuong L. Mai, Kristin Zelley, Valen E. Johnson, Danielle Braun, Kim E. Nichols, Judy E. Garber, Sharon A. Savage, Louise C. Strong, Wenyi Wang

## Abstract

*De novo* mutations (DNMs) are increasingly recognized as rare disease causal factors. Identifying DNM carriers will allow researchers to study the likely distinct molecular mechanisms of DNMs. We developed Famdenovo to predict DNM status (DNM or familial mutation (FM)) of deleterious autosomal dominant germline mutations for any syndrome. We introduce Famdenovo.TP53 for Li-Fraumeni syndrome (LFS) and analyze 324 LFS family pedigrees from four US cohorts: a validation set of 186 pedigrees and a discovery set of 138 pedigrees. The concordance index for Famdenovo.TP53 prediction was 0.95 (95% CI: [0.92, 0.98]). Forty individuals (95% CI: [30, 50]) were predicted as DNM carriers, increasing the total number from 42 to 82. We compared clinical and biological features of FM versus DNM carriers: 1) cancer and mutation spectra along with parental ages were similarly distributed; 2) ascertainment criteria like early-onset breast cancer (age 20 to 35 years) provides a condition for an unbiased estimate of the DNM rate: 48% (23 DNMs versus 25 FMs); 3) hotspot mutation R248W was not observed in DNMs, although it was as prevalent as hotspot mutation R248Q in FMs. Furthermore, we introduce Famdenovo.BRCA for hereditary breast and ovarian cancer syndrome, and apply it to a small set of family data from the Cancer Genetics Network. In summary, we introduce a novel statistical approach to systematically evaluate deleterious DNMs in inherited cancer syndromes. Our approach may serve as a foundation for future studies evaluating how new deleterious mutations can be established in the germline, such as those in *TP53*.

## INTRODUCTION

*De novo* mutations (DNMs) are defined as germline variants (mutations) occurring in the carrier, but not in either of the carrier’s parents (Rahbari et al. 2016). In humans, dozens of DNMs (Conrad et al. 2006; Kondrashov 2003; Kong et al. 2012; Lipson et al. 2015; Lynch 2010; Nachman and Crowell 2000) occur throughout the genome of each newborn. DNMs are considered a major cause for the occurrence of rare, early-onset reproductively lethal diseases (Acuna-Hidalgo et al. 2016). Initial analyses have shown that the rate of germline DNMs increases with paternal (Kong et al. 2012) and maternal age (Wong et al. 2016); plus, DNMs are correlated to a number of factors (Battle and Montgomery 2014), including the time of replication (Francioli et al. 2015), the rate of recombination (Francioli et al. 2015), GC content (Michaelson et al. 2012), and DNA hyper-sensitivity (Michaelson et al. 2012). DNM carriers and their associated factors have always been a focus of genetics research because of their crucial role in the evolution of species (Goldmann et al. 2019).

Not only are DNMs the raw materials of evolution, but they may also shape and determine disease susceptibility (De Ligt et al. 2012; De Rubeis et al. 2014; Iossifov et al. 2014; Krumm et al. 2015; Robinson et al. 2014). DNMs have been recognized as causal factors for rare diseases, such as retinoblastoma and neurofibromatosis, where the *de novo* rates are as high as ∼90% (Dryja et al. 1997) and ∼50% (Evans et al. 2010; Jett and Friedman 2010), respectively. Li-Fraumeni syndrome (LFS) is an autosomal-dominant cancer predisposition syndrome that commonly manifests as soft tissue and bone sarcomas, breast cancer, brain tumors, leukemia, and adrenal cortical carcinomas (Olivier et al. 2003; Nichols et al. 2001) and is associated with germline mutations in the *TP53* tumor suppressor gene (Malkin et al. 1990; Correa 2016), of which 7% - 20% are *de novo* (Gonzalez et al. 2009). Clinically, it is of interest to identify *de novo* carriers of a *TP53* germline mutation, who may otherwise go unnoticed until they or their offspring develop multiple cancers, to provide them with information on current standard LFS screening and surveillance protocols. Understanding the cause of *de novo* mutations in cancer genes like *TP53* will help develop strategies for early identification, which will have a novel and significant impact on the field of human genetics.

Predicting a DNM in a gene is not trivial when the parental mutation statuses are unknown. Incomplete family history may skew observational criteria and the prediction may suffer from low sensitivity and specificity. Mendelian risk prediction models (Chen et al. 2004) have been used successfully for accurate prediction of deleterious mutation status in familial breast and ovarian (Euhus et al. 2002), bowel (Chen et al. 2006b), and pancreatic (Wang et al. 2007) cancers; melanoma (Wang et al. 2010); and most recently in LFS (Peng et al. 2017; Renaux-Petel et al. 2018). We developed a novel statistical approach, Famdenovo, to predict a mutation carrier’s DNM status, i.e., whether the carrier’s deleterious mutation was due to familial inheritance or alternatively arose from a new mutation in that individual. Built on the principles of Mendelian models, Famdenovo predicts a new biological outcome, DNM status. For each particular set of gene(s) of interest, three sets of input parameters need to be estimated from a population study in order to build Famdenovo: the penetrance, the allele frequency, and the *de novo* mutation rate (percent of *de novo* mutations among all germline mutations) of the deleterious alleles.

DNMs are rare, and therefore require a large amount of high-quality data to be studied thoroughly. We introduce Famdenovo.TP53, which was modeled using input parameters estimated from published studies and validated on a large cohort consisting of 324 LFS family cohorts that were collected from four major cancer centers in the US, with *TP53* genetic testing results available in at least one trio per family in 186 families. The computer-based and data-driven identification of deleterious DNM carriers in *TP53*, who are otherwise hidden in a wide population, enabled a new perspective of DNMs that is gene and disease-centric. In addition, we introduce Famdenovo.BRCA for hereditary breast and ovarian cancer syndrome (HBOC), and apply it to a small set of family data from the Cancer Genetics Network (CGN).

## RESULTS

### Famdenovo for predicting the deleterious DNM status among mutation carriers

Famdenovo is based on widely-used Mendelian models (Chen et al. 2004, 2006b; Parmigiani et al. 1998; Wang et al. 2007, 2010) and calculates the probability that a deleterious germline mutation is *de novo*. The *de novo* probability of interest can be written as the Pr(father is a noncarrier, mother is a noncarrier | child is a mutation carrier, family history). We calculate this probability as a function, g(*c, f, m*), of pedigree-based carrier probabilities for the child (*c*), father (*f*) and mother (*m*), respectively. We derived the g() function using Bayes’ rule (as detailed in the Methods section), and obtained input values using Mendelian models as the corresponding marginal probabilities for one individual’s *c, f*, and *m*, respectively. Each of the marginal probabilities is calculated using the Elston-Stewart algorithm (Fernando et al. 1993). Famdenovo then applies Bayes’ rules to estimate *de novo* probabilities in deleterious mutations. Similar to previously developed Mendelian models, Famdenovo can be adapted for specific disease-gene associations by setting its input parameters accordingly: the penetrance of the disease-gene, the allele frequency of the gene(s), and the *de novo* mutation rate. We have incorporated well-validated parameter estimates in Famdenovo.TP53 for LFS-*TP53* (Peng et al. 2017) and Famdenovo.BRCA for HBOC-*BRCA1/2* (Chen et al. 2006a; Parmigiani et al. 2007).

**Figure 1** depicts three scenarios of a hypothetical pedigree resulting in different DNM probabilities in *TP53* given by Famdenovo.TP53. **Figure 1A** shows a pedigree with four cancer cases: two soft tissue sarcomas (STS) at 25 and 27 years old, one breast cancer at 40 years old, and another STS at 20 years old. The arrow points to the counselee who is a *TP53* mutation carrier, and the probability of the mutation being *de novo* for the counselee is 0.0033. It is likely that this is a familial mutation inherited from the counselee’s mother. **Figure 1B** shows a similar pedigree with three cancer patients, with one less affected relative in the maternal branch of the counselee compared with **Figure 1A**, and with an older age of diagnosis for the counselee’s mother, which increases the predicted probability of the mutation being *de novo*. The *de novo* probability given by Famdenovo.TP53 is 0.47. **Figure 1C** shows a similar pedigree with only two cancer cases in the counselee and counselee’s offspring, compared with **Figure 1B**. Because there are no longer any cancer patients in the older generations and the counselee’s siblings are all healthy, it is most likely that this mutation is *de novo*. Famdenovo.TP53 predicted this mutation to be a DNM, based on the .89 probability of the mutation being *de novo*.

**Figure 1.**
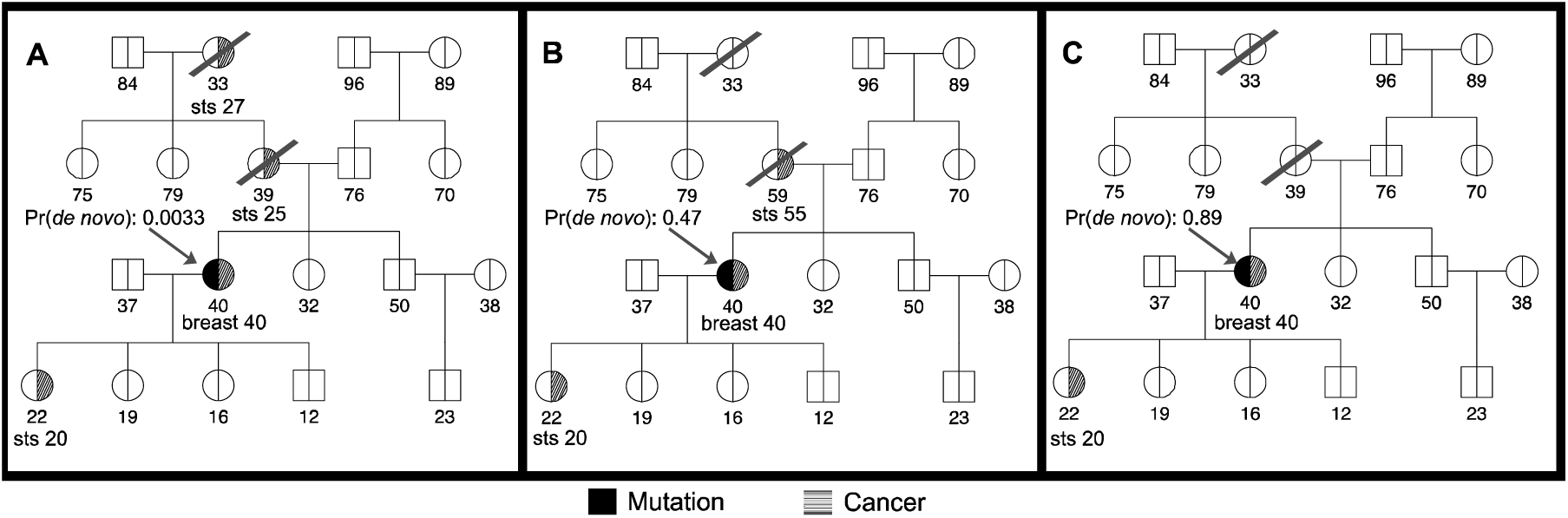
Illustration of clinical counseling using Famdenovo based on family history. The number below each individual is the age at last contact (healthy or affected with cancer). The arrow points to the counselee who is known to be a *TP53* mutation carrier. The *de novo* probabilities shown are given by *Famdenovo*.*TP53*. **(A)** A pedigree with four cancer patients. **(B)** A pedigree with three cancer patients. **(C)** A pedigree with two cancer patients.

### Validation of Famdenovo.TP53 in individuals with known DNM status in *TP53*

We evaluated the performance of Famdenovo.TP53 based on germline *TP53* tested individuals with known DNM status from the four cohorts. Among the 324 LFS family pedigrees we collected, 186 families had a known DNM status and 138 had an unknown DNM status (**Figure 2, Table S1**). **Table 1** provides a summary of the demographics for all four study cohorts. The largest set of DNM carriers was contributed by the MD Anderson (MDA) cohort. In total, there were 42 known DNM carriers and 144 families with known FM carriers.

**Table 1.** Summary of demographic information for all the families with *TP53* mutation carriers from the four cohorts.

| Cohort |  |  | Individuals per families |  | Age at first diagnosis |  |
| --- | --- | --- | --- | --- | --- | --- |
| MDA | No. of Families | 140 | Mean | 45 | Mean | 42.3 |
|  | Max Family Size | 151 | Standard Deviation | 31 | Standard Deviation | 22.2 |
|  | Min Family Size | 4 |  |  |  |  |
| NCI | No. of Families | 78 | Mean | 24 | Mean | 33.9 |
|  | Max Family Size | 130 | Standard Deviation | 17 | Standard Deviation | 22.1 |
|  | Min Family Size | 3 |  |  |  |  |
| DFCI | No. of Families | 91 | Mean | 34 | Mean | 38.9 |
|  | Max Family Size | 107 | Standard Deviation | 25 | Standard Deviation | 20.5 |
|  | Min Family Size | 3 |  |  |  |  |
| CHOP | No. of Families | 15 | Mean | 26 | Mean | 43.4 |
|  | Max Family Size | 45 | Standard Deviation | 12 | Standard Deviation | 25.2 |
|  | Min Family Size | 13 |  |  |  |  |

**Figure 2.**
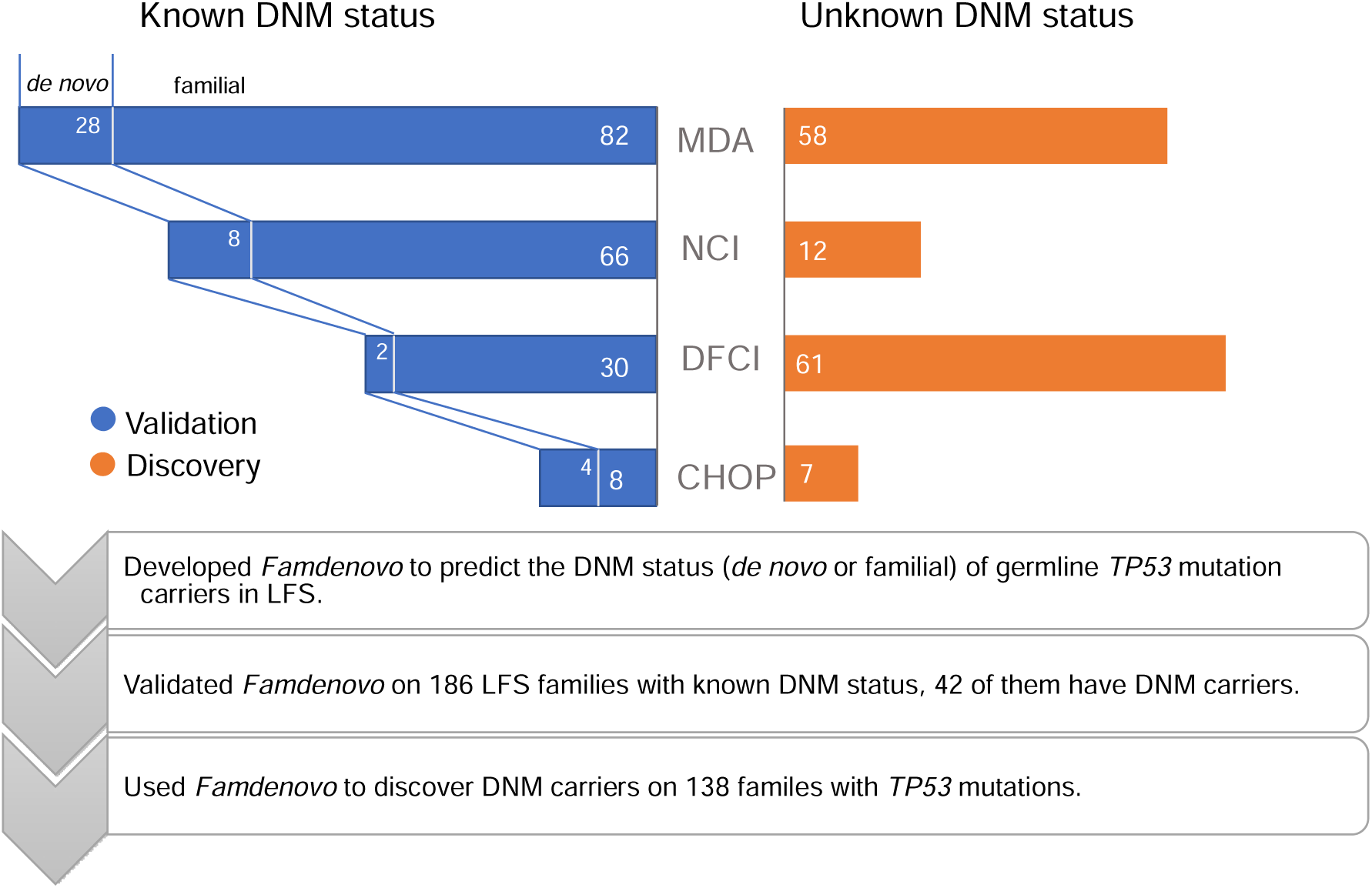
Study design and methods. The study included four cohorts: MD Anderson (MDA), National Cancer Institute (NCI), Dana-Farber Cancer Institute (DFCI), and Children’s Hospital of Philadelphia (CHOP) in our study. A subset of families within each cohort have mutation testing results from the parents, and therefore the true DNM status is known. These families form the validation set of our study. In the other families, the family members’ DNM statuses are unknown, and they form the discovery set.

We used observed:estimated ratios (OEs) and receiver operating characteristic (ROC) curves to evaluate the calibration and discrimination of our model. **Figure 3** shows the ROC curves for the validation set from the four cohorts combined. The total sample size for the validation was 186 families; the corresponding area under the curve (AUC, or concordance index) was 0.95 (95% CI: [0.92, 0.98]), which demonstrated that Famdenovo.TP53 performed well in discriminating DNM carriers from FM carriers. The calibration was satisfactory, with some under-prediction and an OE of 1.33 (95% CI: [1.09, 1.63]).

**Figure 3.**
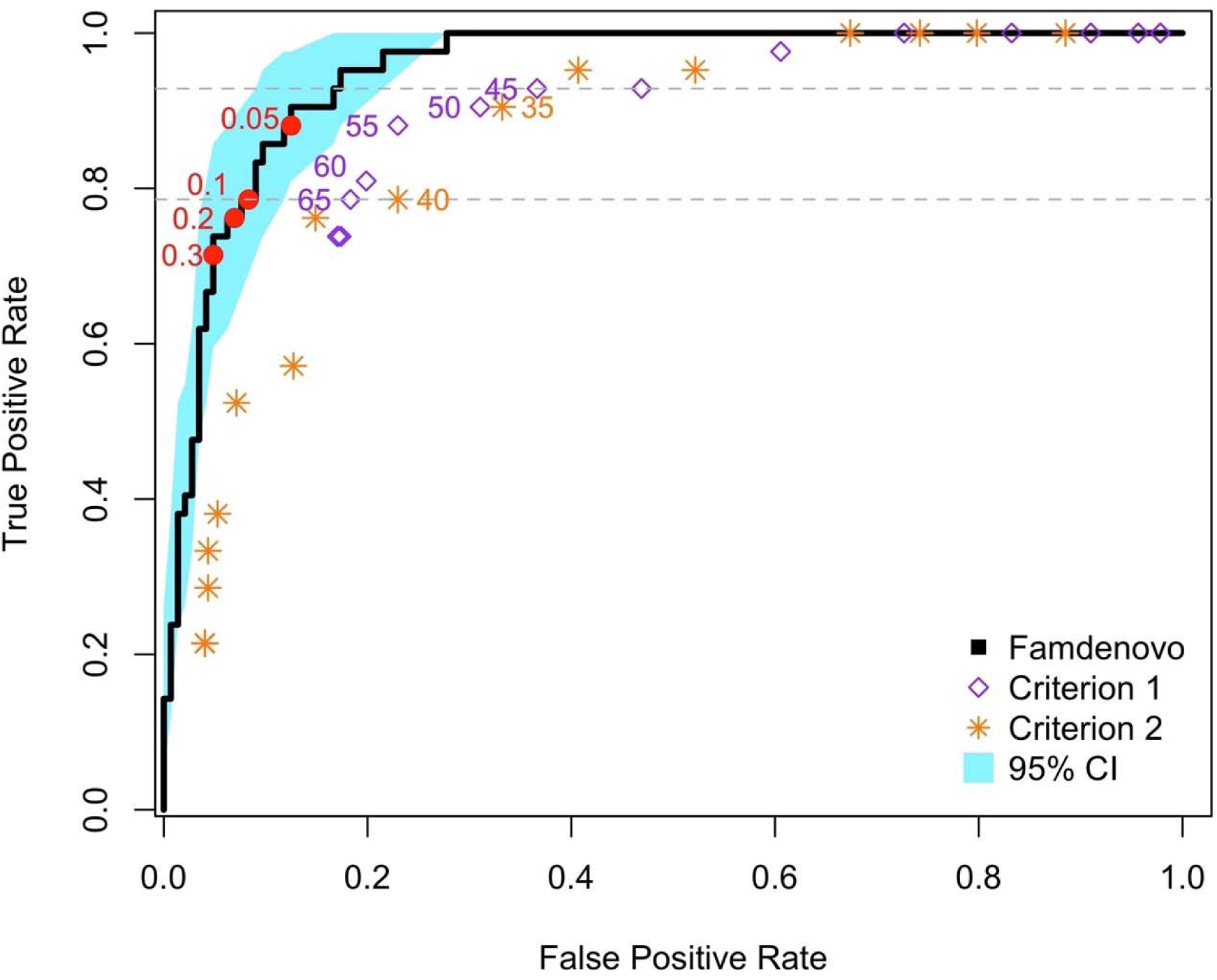
ROC curves for the validation of Famdenovo.TP53. The x axis is the false positive rate (i.e., 1 – specificity) and the y axis is the true positive rate (i.e., sensitivity). The black curve corresponds to the results of Famdenovo.TP53 on the 186 families in the validation set and the blue shade shows the 95% confidence interval. The red circles represent the different *de novo* cutoff probabilities for the model. The purple diamonds correspond to the results of criterion 1 at a cut-off age (provided next to the diamonds). The orange stars represent the result of criterion 2 at a cut-off age (provided next to the stars). The gray dotted lines show the range of cut-off ages, 45 to 65, where a partial AUC is calculated for criterion 1.

We also compared our method with two simple criterion-based prediction models that do not require the collection of extended pedigree data (**Table 2**). Criterion 1 predicts a mutation to be *de novo* if the mutation carrier does not have a parent diagnosed with cancer before a cut-off age, e.g., 45 or 60. Criterion 2 predicts a mutation to be *de novo* if, in addition to criterion 1, neither of the grandparents was diagnosed with cancer before the cut-off age. Criterion 1 is more relaxed than criterion 2. **Figure 3** shows that both criteria (over a range of cut-off ages) fall outside of the 95% confidence interval band of our method. Neither of the criteria is able to reach the full range of the ROC curve. Criterion 2 follows the same trend as criterion 1. Moreover, for each criterion we calculated a partial AUC area with cutoff ages from 45 to 65, which were then rescaled to the full range of 0 to 1. The rescaled AUC for criterion 1, the better performing of the two criteria, is still much lower than Famdenovo.TP53, with a difference of 0.14 (95% CI: [0.054, 0.20], **Table 2**). Our results demonstrate that Famdenovo.TP53 outperforms criterion-based prediction models, and that extended pedigrees with cancer diagnosis information are needed for the accurate identification of DNM carriers.

**Table 2.** Comparison of AUCs of Famdenovo.TP53 and various criterion-based prediction models, where 95% CI were obtained by bootstrapping.

| AUC | Mean | 95% CI |
| --- | --- | --- |
| Famdenovo.TP53 | 0.95 | [0.92, 0.98] |
| Criterion 1 | 0.82 | [0.78, 0.87] |
| Criterion 2 | 0.80 | [0.77, 0.84] |
| Famdenovo -<br>criterion 1 | 0.14 | [0.054, 0.20] |
| Famdenovo -<br>criterion 2 | 0.15 | [0.087, 0.20] |

We evaluated the sensitivity and specificity of Famdenovo.TP53 under various prediction cutoff values using the validation set (**Table S2**) in order to identify an optimal classification cutoff for this dataset. If the predicted probability was equal to or above the prediction cutoff, the individual was classified as a DNM carrier. As the cutoff probability increases, the sensitivity decreases and specificity increases. We chose a cutoff of 0.2, which provided a good tradeoff of a sensitivity of 0.76 and a specificity of 0.93. Maintaining a high specificity is essential to allow for downstream clinical interpretation of *de novo* mutations. Using the observed deleterious DNMs in our *TP53* clinical cohorts, we estimated that Famdenovo.TP53, at a cutoff=0.2, will achieve a positive predictive value (PPV) of 0.78 and a negative predictive value (NPV) of 0.92. Without using pedigree information and model-based analysis like Famdenovo.TP53, criterion 1 will only achieve a PPV of 0.63 and an NPV of 0.92 (cutoff age = 65, sensitivity = 0.8, and specificity = 0.8).

We also conducted a sensitivity analysis to evaluate how the pedigree size affects the validation results. The range of pedigree sizes for each of the four cohorts is listed in **Table 1** (see also **Figure S1**). We divided the 186 families in the validation dataset into two groups: large families and small families, using the median size of all families (n = 29) as the cut-off size. We obtained similar results under the two scenarios (small families: AUC 0.96, 95% CI [0.91, 0.99], large families: AUC 0.94, 95% CI [0.89, 0.98]). These results support the robustness of our model by showing that Famdenovo is capable of accurately predicting *de novo TP53* mutations in a range of family sizes.

We further conducted a sensitivity analysis on the two input parameters: the prior values for the deleterious allele frequency and the DNM rate. These values are expected to be dominated by family history data that provide strong information during the calculation of posterior probabilities. Tuning the two parameters barely influenced the AUC or OE (**Table S3**).

### Discovery in individuals with unknown DNM status using Famdenovo

We applied Famdenovo.TP53 to the 138 families with unknown DNM status in the discovery set from the same four cohorts. **Figure 4** provides the number and percentage of predicted DNM carriers at a cutoff of 0.2. In total, 40 (95% CI: [30, 50]) out of the 138 families were predicted to be *de novo*. The number of predicted *de novo* families for MDA, NCI, DFCI, and CHOP were 23 out of 58 (95% CI: [16, 30]), 7 out of 12 (95% CI: [4, 10]), 7 out of 61 (95% CI: [3, 12]), and 3 out of 7 (95% CI: [1, 5]), respectively. MDA and CHOP had higher percentages of DNMs, likely due to differences in ascertainment across cohorts. Sensitivity analyses using cutoffs from 0.05 to 0.3 gave consistent DNM results (**Table S4**).

**Figure 4.**
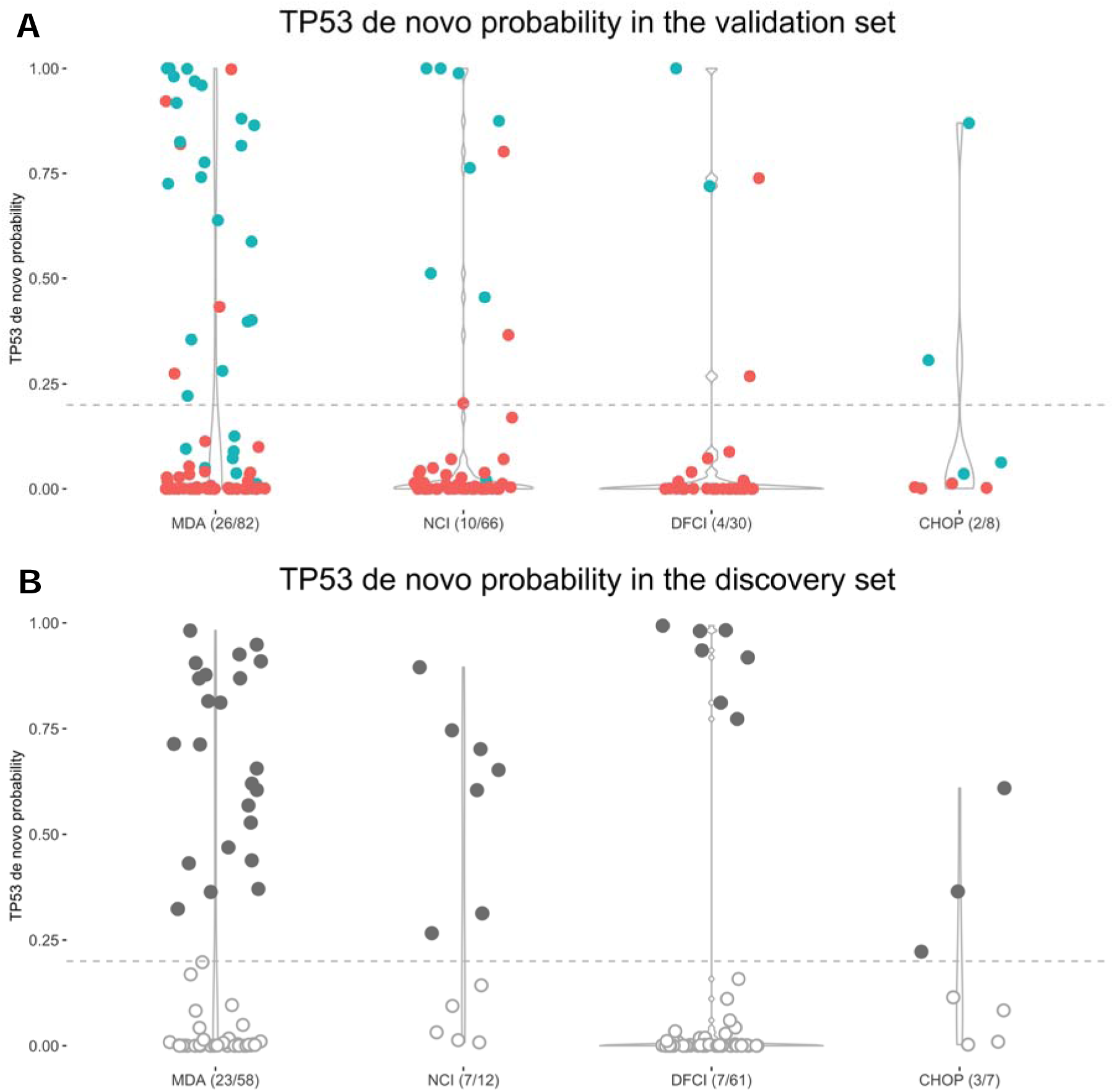
Distributions of the *de novo* probabilities in validation and discovery sets in the four cohorts. The y axis is the predicted *de novo* probability by Famdenovo.TP53. The width of each column changes over the y axis, representing the percent of individuals who have the corresponding range of probability values. The individuals with points above the gray line (cutoff = 0.2) were predicted to be DNM carriers. The x axis gives the names of the cohorts and the corresponding predicted numbers of DNM carriers out of the total number of individuals. (**A**) In the validation set, the individuals with blue dots were true DNM carriers, and those with red dots were true FM carriers. (**B**) In the discovery set, the individuals with solid gray dots were predicted to be DNM carriers, and those with empty dots were predicted to be FM carriers.

### Association of deleterious DNMs in *TP53* with clinical outcomes

Combining the validation and discovery sets resulted in a total of 82 DNMs from 82 families and 450 FMs from 242 families, which equipped our study with a sample size that was meaningful to evaluate associations between mutation status and a diverse range of individual outcomes: sex, cancer types, multiple primary cancers, age of cancer diagnosis, parental ages, and types of mutation.

Comparing DNM to FM carriers in 397 cancer patients (**Table S5**), we observed more females than males (Odds Ratio (OR) = 1.78, 95% CI: [1.01, 3.13]). Considering the strong association between sex and breast cancer, we asked whether this significant finding comes from ascertainment bias. We performed another test within the probands, i.e., the index person that brings the family into the research cohort. This time, in contrast, we found no sex differences (34% DNMs in female and 32% DNMs in male, OR=1.08, 95% CI: [0.58, 2.03]). All but two DNM carriers were probands. Therefore, we infer the former significant finding was due to an imbalanced addition of FM carriers, who were relatives of the probands, to the female and male categories, i.e., a numerical artifact. In the following association analyses, we present comparisons within the probands only.

Among 233 probands with cancer information (**Table S6**), we observed an OR = 1.9 (95% CI: [1.09, 3.33]) for DNMs in breast cancer as compared to other cancer types. We further divided the probands into three age categories: [0, 20), [20, 35) and 35+, using age of last contact. We observed, within the [20, 35) age category, an OR = 5.98 (95% CI: [1.81, 19.76]) for DNMs in breast cancer as compared to other cancer types, whereas a similar test in the age 35+ category was not statistically significant. This association is again due to ascertainment of DNMs through patients with early-onset breast cancer and not with other cancer types (see **Table S7** for commonly used ascertainment criteria (Li and Fraumeni Jr. 1969; Tinat et al. 2009)). Within the [20, 35) age category and probands with breast cancer, where the ascertainment of each individual is consistent, we found 23 DNM carriers (29% of all proband DNM carriers) and 25 FM carriers (16% of all proband FM carriers), resulting in a 48% (sd = 7.2%) DNM rate. This high proportion of DNMs is also observed within the validation set only (n = 11 vs. 8, **Table S6C**). Sensitivity analyses using various thresholds of age of last contact showed consistent results (**Table S6D**), supporting the robustness of this finding.

We evaluated the association between the DNM status and the risk of developing subsequent multiple primary cancers (MPC) within the probands. **Table S8A** shows that more patients with DNMs presented multiple primary than single primary cancers (OR = 1.77, 95% CI: [1.01, 3.11]). Similarly, we attribute such association to the ascertainment practice, i.e., ascertaining patients with a personal history of multiple primary cancers (at least one cancer is at an early onset) is part of the clinical criteria used for patient ascertainment in our study cohorts (Li and Fraumeni Jr. 1969; Tinat et al. 2009). In an attempt to adjust for ascertainment to make the DNM and FM carriers comparable, we looked within the [20, 35) age category at probands with MPC. We found 17 DNM carriers and 22 FM carriers, resulting in 44% of DNMs within this combined cohort population (sd = 7.9%, **Table S8B**). This high proportion is also observed with the validation set (n = 10 vs. 8, **Table S8C**). This result is congruent with the previous observation of a high DNM rate in probands with early-onset breast cancer.

**Figure 5A** shows the distributions of cancer diagnosis age as stratified by cancer type and sex for the combined set. We observed DNMs in all of the eight LFS spectrum cancers: breast, brain, leukemia, choroid, adrenocortical carcinoma, osteosarcoma, soft tissue sarcoma, and lung, as well as other invasive cancers that are not within the LFS spectrum. Other than breast cancer, most cancer types have both female and male DNM carriers, with the exceptions of lung cancer and leukemia. For lung cancer, DNMs were only presented in females (n = 5, mutation type: NC_000017.10:g.7675232G>A (p.S127F), NC_000017.11:g.7675122G>A (p.P152L), NC_000017.11:g.7674887_7674889delinsTGG (p.R213W), NC_000017.11:g.7675190C>T (p.G245S), NC_000017.11:g.7670699C>T (p.R337H), **Table S9**) and not males. For leukemia, DNMs also were only presented in females (n = 2, mutation type: NC_000017.11:g.7673776G>A (p.R282W), **Table S9**). We found no difference in ages of diagnosis between DNM and FM carriers across cancer types, including breast cancer (**Figure 5A**).

**Figure 5.**
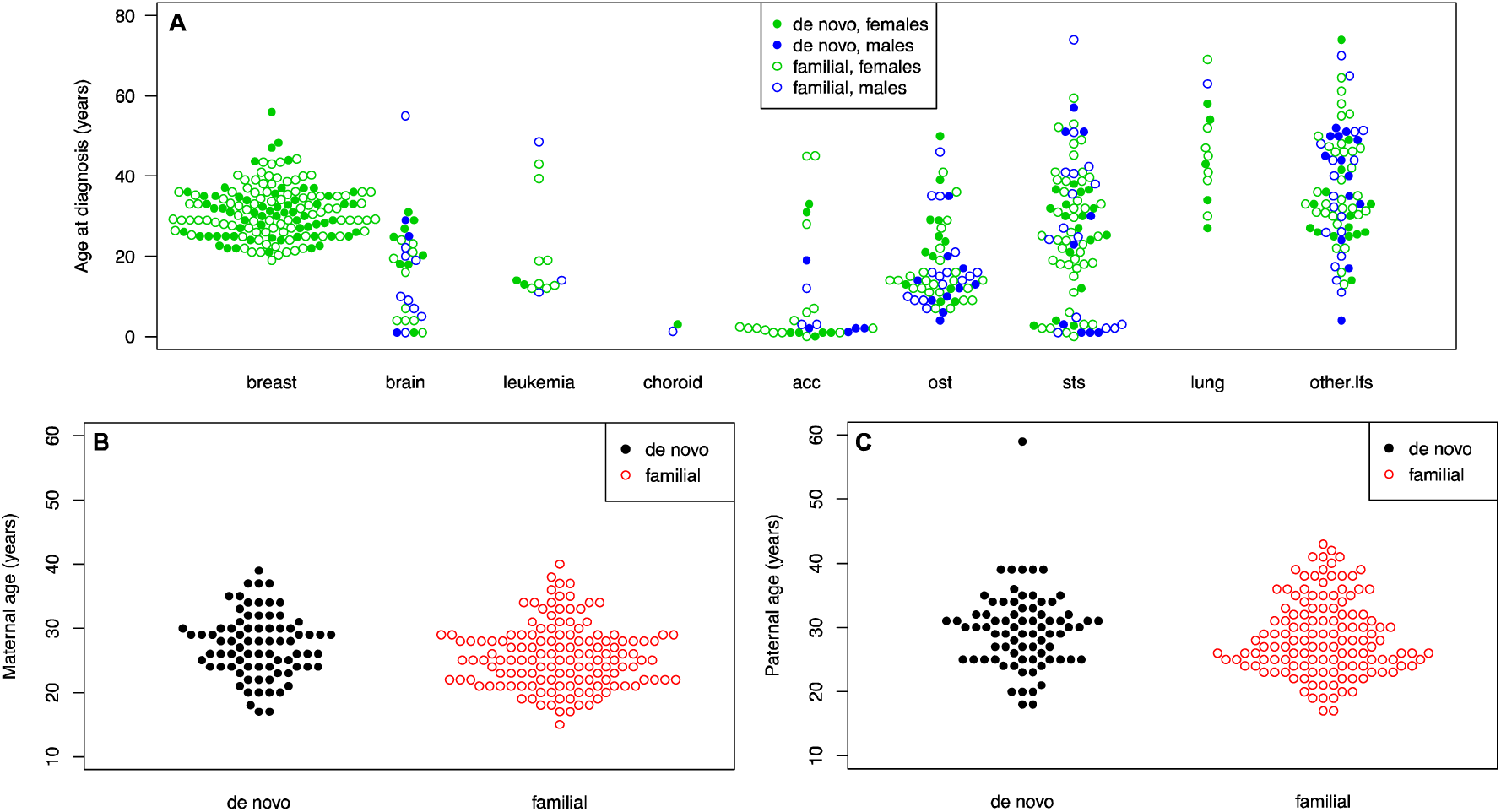
Ages of cancer diagnosis and parental ages in DNM and FM carriers. **(A)** Ages of cancer diagnosis for all patients. The x axis represents types of different cancer types. Cancers were classified by site: LFS spectrum (osteosarcomas, soft-tissue sarcomas, breast, brain, adrenal, lung cancers, and leukemia) and non-LFS spectrum (prostate, colon, kidney, or thyroid cancers, and others). The green dots are for females, while the blue dots are for males. **(B)** Distribution of paternal ages. **(C)** Distribution of maternal ages. All solid dots represent the DNM carriers.

We also investigated whether the parental ages (paternal or maternal) were different among *TP53* DNM and FM carriers. Parental age refers to the age of the parent at the time of child birth. Literature on the origins of DNMs suggested their occurrence rate across the genome was correlated with paternal age (Kong et al. 2012). Little is known about occurrence of deleterious DNMs within a given gene with regard to parental ages. Here parental age is obtained by calculating the difference between the birth year of the child and the parent. **Figures 5B and 5C** show the distributions for the combined set. We observed that both paternal age (mean = 29, sd = 5.8) and maternal age (mean = 27, sd = 5.0) were similarly distributed between DNMs and FM carriers. This is consistent with what was reported recently in another study (Renaux-Petel et al. 2018) for deleterious DNMs within *TP53*.

**Table S10** summarizes the frequencies of DNMs in each of the four cohorts MDA, NCI, DFCI, and CHOP within the validation and discovery sets. In this study of *TP53* mutations, there were a total of 49 variants that were recorded in DNMs (**Table S9**), and a total of 82 variants recorded in FMs. **Figures S2** summarizes the occurrences of DNM and FM variants in the validation and discovery sets, as well as their overlap in mutation types (24.5% and 34.1%). Multiple hotspot mutations were presented in FMs. Among them, two *TP53* hotspot mutations (Walerych et al. 2012), NC_000017.11:g.7674220C>T (p.R248Q) and NC_000017.11:g.7673802C>T (p.R273H), were the most enriched in DNMs (p.R248Q, n = 4, cancer types: adrenocortical carcinoma, breast, osteosarcoma, soft tissue sarcoma, **Table S10**). In particular, NC_000017.11:g.7674221G>A (p.R248W) mutations were missing among the DNMs list for this study. Among the FMs, p.R248W was as prevalent as p.R248Q (n = 7 vs. 9 out of 242). The observation of n = 0 out of 82 DNMs for p.R248W was marginally significant (p-value = 0.09, based on a Poisson distribution). The cancer types of FM-p.R248W carriers encompass the cancer spectrum of LFS, including breast cancers. The frequencies of DNMs at different amino acid positions (**Figure 6**) did not differ significantly in the validation and discovery sets, supporting the validity of predicted DNMs in the discovery set.

**Figure 6.**
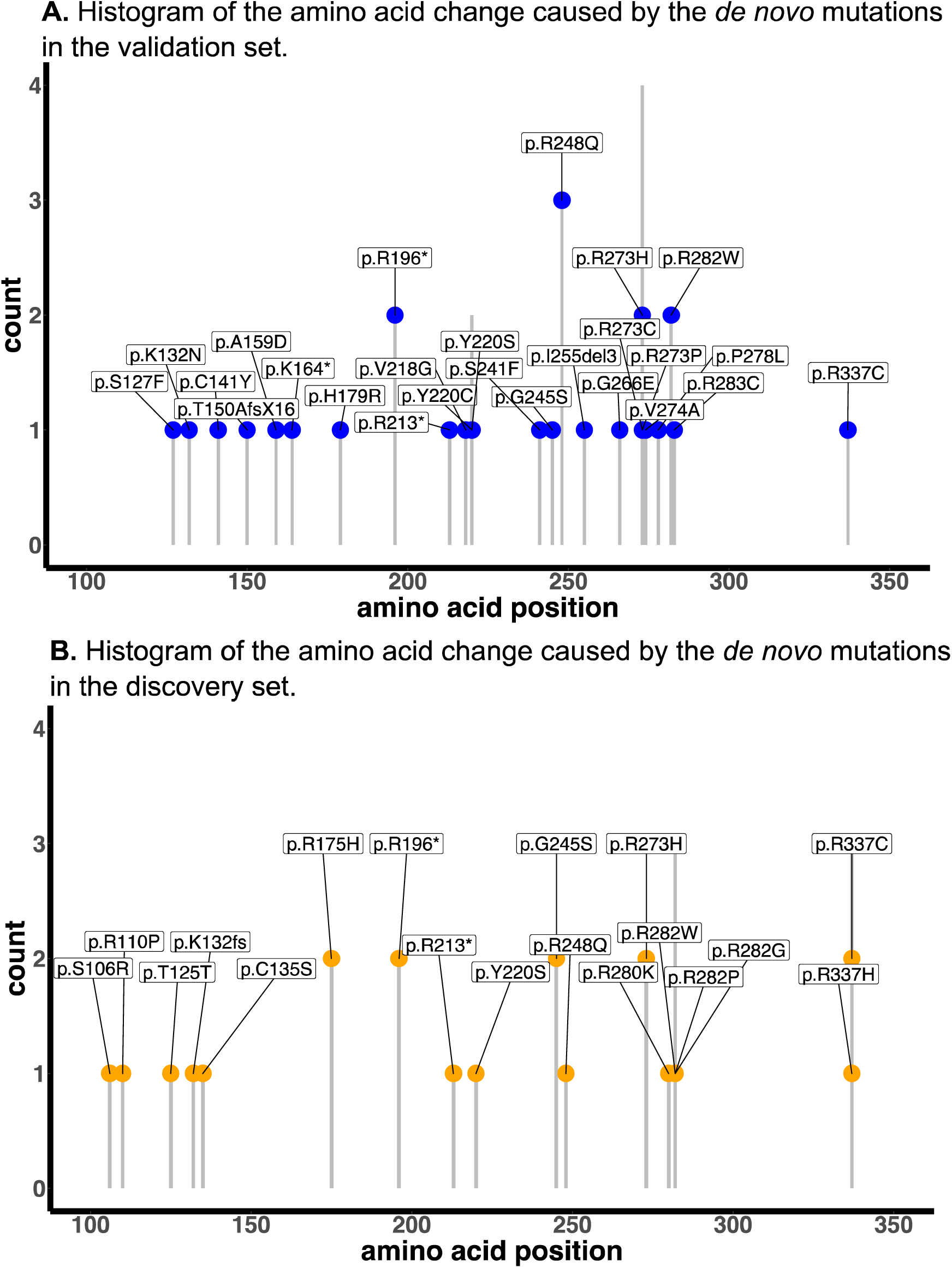
Histograms of DNM frequency at different amino acid positions in the validation and discovery sets. (**A**) The validation set. (**B**) The discovery set. The y axis shows the count of DNMs. The x axis shows the amino acid positions.

### Deleterious DNM status among mutation carriers in *BRCA1/2*

We used a dataset with 39 extended pedigrees from the Cancer Genetics Network (Anton-Culver et al. 2003): 23 families had *BRCA1* mutation carriers, and 16 families had *BRCA2* mutation carriers (**Table S11**). Among them, we predicted a total of 7 (18%) DMNs using a cutoff of 0.15. Among the 7 predicted DMNs, 3 people (13%) were *BRCA1* mutation carriers, and 4 people (25%) were *BRCA2* mutation carriers (**Figure S3**). Visual inspection of the family cancer history of these 7 families (**Figure S4**) supports the validity of the Famdenovo predictions. The median probability of the 59 predicted FM carriers is 0.005. Due to a lack of genetic testing information of both parents, we were limited to positive testing results from some parents or siblings, which provided us with only true negatives for validation. A total of 38 people from 15 families were confirmed through parent-child relationship, or inferred to be FMs in the case of having multiple siblings tested positive. Famdenovo.BRCA also predicted all of them as FMs (specificity = 100%), which demonstrated the accuracy of Famdenovo.BRCA.

## DISCUSSION

Deleterious germline DNMs are more prevalent than previously thought. For cancer genes such as *TP53*, we estimated a ∼1:1 ratio of deleterious DNMs compared to FMs in our LFS cohorts, supporting the importance of studying its etiology. To enable future genomic studies, we have developed a statistical method and tool, Famdenovo, to predict the deleterious DNM status in cancer genes using family history data. Famdenovo.TP53 showed excellent performance, an AUC of 0.95, in discriminating DNM carriers from FM carriers of *TP53* deleterious germline mutations in 186 families collected from four clinical cohorts with varying ascertainment criteria. In our case study using 324 germline *TP53* pedigrees, Famdenovo.TP53 identified a total of 82 deleterious DNM carriers (increased from only 42 using trio-based genetic testing results, 80 of which were cancer patients) and 450 deleterious FM carriers (317 of which were cancer patients), contributing to a sample size that is large enough to test for associations with patient outcomes: sex, cancer types (>9 different types), multiple primary cancers, age of cancer diagnosis, parental ages, and mutation types. We provide a reference of cancer spectrum and mutation spectrum to contrast *TP53* DNM and FM carriers as a resource for future studies, complementing the ongoing efforts to accurately annotate *TP53* variants (Leroy et al. 2017; Tikkanen et al. 2018). Hotspot mutations p.R248Q and p.R273H, the most commonly mutated residues in breast cancer (Walerych et al. 2012), were the most frequently observed mutation types among DNMs. In contrast, p.R248W was not observed in DNMs, but was prevalent in FMs.

The observed discrepancy in frequency of p.R248W in DNMs versus FMs, but not in p.R248Q, is consistent with the literature on functional differences in the two mutations (Walerych et al. 2012). When mutated in cancer cells, p.R248W has recently been reported to induce a much stronger response from T cells than p.R248Q (Malekzadeh et al. 2019). Germ cells containing DNMs are sustained in a microenvironment consisting of parental cells that do not carry such DNMs, hence mimicking heterogeneous cell-cell interactions that are similar to those presenting somatic mutations. In our data, it is plausible that a neoantigen-like response to p.R248W mutated germ cells has helped destroy these cells so that they are not viable for reproduction. If these germ cells have survived however, then their offspring may no longer incur such response, as all cells now contain this mutation. This hypothesis is supported by our observation of a high prevalence of p.R248W in FMs. An alternative hypothesis to explain this observation is that DNMs in p.R248W were simply not ascertained by this study, for unknown reasons, as discussed further below.

DNM carriers do not manifest via family history data and are likely to be identified in clinic only when they have developed multiple cancers, very early-onset breast cancer, or their offspring have developed cancer. They are hence currently under-represented in cancer genetics studies. We estimate that our four study cohorts may miss up to 45% of deleterious DNMs in *TP53*, due to the lack of genetic testing for cancer patients with no family history or healthy individuals who carry these mutations. Correspondingly, we estimate a population prevalence of 0.00076 for deleterious *TP53* mutations, which is 26% higher than shown in previous studies (Peng et al. 2017). Our finding, that deleterious DNMs of *TP53* are as frequent as FMs in early-onset breast cancer provides strong evidence to support the NCCN guideline for women with early onset breast cancer to be offered testing for *TP53*, regardless of family history, to capture the DNM carriers who are not as rare as previously thought. Our results provide a quantitative evaluation on DNMs to genetic counselors so that they may tailor sessions to carefully explain the new knowledge. More importantly, DNMs are sparse in the general population although their downstream effect is detrimental for families. Among the four cohorts, there were 138/324 probands with missing parental genotypes. In these cases, Famdenovo.TP53 would be useful to predict DNMs. Identification of patients with DNMs in order to study them collectively will further our understanding of the molecular mechanism underlying deleterious DNMs, and it is essential for future identification of the hidden DNMs for genetic testing.

Missing data in parental genotypes is more prevalent in HBOC cohorts, which restricts us from conducting an extensive validation of Famdenovo.BRCA. Genetic counselors suggest that it may be due to the higher age of BRCA1/2mutation carriers at the time they enter a study, resulting in reduced parental follow-up compared to TP53 mutation carriers (commonly found in childhood cancer patients).

We remain cautious with the downstream interpretations of Famdenovo predicted DNMs and, to a larger extent, to any other biological factors that may confound our interpretation. We enumerate the following five considerations in the context of *TP53*, which are applicable to *BRCA1/2* and other future genes of interest. First, mosaicism in parents could be partially contributing to DNMs in the children (Renaux-Petel et al. 2018). If a parent presents an LFS-like phenotype due to a mosaic *TP53* mutation, Famdenovo.TP53 would consider the parent as a germline mutation carrier, hence deducing the child to be an FM carrier. The DNM status of a child under this condition would then be missed by Famdenovo.TP53. Such a condition is rare, at ∼5% (Renaux-Petel et al. 2018), and of less concern to clinical practice, as this population is already targeted by genetic testing. Patients with an LFS-like phenotype are often tested, first with Sanger sequencing, then with NGS at high depth, until mosaicism is confirmed. Second, somatic clonal expansion in cancer patients may create a look-alike germline *TP53* mutation, resulting in a false positive in DNM status. However, such observation was reported in a general panel-testing population, which is supposedly >99.9% without *TP53* mutations, hence presenting an expected low positive predictive value (Weitzel et al. 2018). By focusing on clinically ascertained families, we do not expect aberrant clonal expansion to be the cause of the identified DNMs. Furthermore, we only observed two DNM carriers who have ever had leukemia, a cancer type known to present clonal hematopoiesis. Third, Famdenovo.TP53 assumes the mutations are deleterious and their effect on cancer outcomes follow the penetrance curves previously estimated from a set of families. Variants with unknown significance in *TP53* were excluded from the study. Fourth, DNMs across the genome were reported to be shared in siblings (Jónsson et al. 2018). Within a single gene like *TP53*, however, we did not observe sibling sharing. Last, cutoffs for DNM probabilities to classify the mutation carriers into DNM and FM carriers were determined on the same dataset. Future validation studies are needed to determine one or several recommended cutoffs for clinical decision-making.

Because of our study design, distinct from existing DNM studies that are genome-wide and across many individuals, we are limited to a small number of mutations, which even with the addition of the Famdenovo*’*s discovery set would not be sufficient to perform sequence context-based analyses that would follow those existing DNM studies. In contrast, what existing studies cannot provide is Famdenovo’s laser-beam view of the impact of *de novo* mutations in a single gene and a single disease syndrome. Both study designs should be carried out in order to provide a well-rounded understanding of *de novo* mutations.

In summary, we present a new epidemiological study design, enabled by a statistical method called Famdenovo, to evaluate the clinical impact of deleterious *de novo* mutations in cancer genes. We introduce both Famdenovo.TP53 and Famdenovo.brca and illustrate both models using family history cohorts. This study design is important for future genetic research, as well as the clinical management of patients and their families as we learn new biology and associated clinical risk for DNM carriers. Famdenovo was developed as a freely available R package and Shiny web app (http://bioinformatics.mdanderson.org/public-software/famdenovo).

## METHODS

### Model development

Famdenovo estimates the probability of *de novo* mutation in any designated family member who carries a deleterious germline mutation in a given gene, based on a detailed family disease history and values of genetic parameters, such as the penetrance and prevalence, for the gene and the corresponding diseases. The prevalence for gene mutations is expressed as *Pr(G)*, where *G* denotes genotype, which could be wildtype (denoted as 0), a heterozygous mutation (denoted as 1), or a homozygous mutation (denoted as 2). *Pr(G)* for three genotypes can be derived from the prevalence of the mutated alleles using the Hardy-Weinberg equilibrium. The penetrance is the probability of developing the disease at a given age for individuals who carry the germline mutations. The disease of interest is an inherited syndrome, such as Li-Fraumeni syndrome or HBOC. The genes (one or more) of interest are those that are known to be the major cause of the corresponding inherited syndrome when they are mutated. The germline mutations here correspond to deleterious mutations that are accepted in clinics as mutations that would categorize the carriers as high risk for developing the disease in the future and who will be suggested to adhere to a series of cancer prevention protocols. Other mutations in these genes, including those that are variants of unknown significance, are considered as wildtype in the model. Either the dominant or recessive mode of inheritance can be assumed in Famdenovo.

Let ***P*** be the pedigree information of the family (e.g., the pedigree drawn in **Figure 1**), let ***D*** be the disease phenotype information of all family members, and *G*_*C*_ be the genotype of the carrier, then the probability of the mutation being *de novo* can be written as Pr(*G*_*c*_ is *de novo* | *G*_*c*_ is germline,**D**,**P**), referred to as the “*de novo* probability.” To calculate Pr(*G*_*c*_ is *de novo* |*G*_*c*_ is germline,**D**,**P**), we have the following equation:

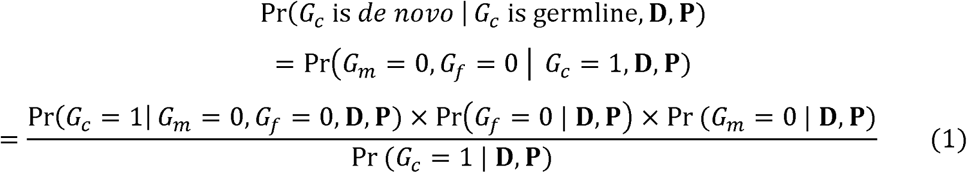

where *G*_*m*_ is the genotype of the mother and *G*_*f*_ is the genotype of the father. In equation (1), all four probabilities are usually difficult to obtain through direct calculations. Hence, we apply Mendelian models to derive them. Let the family cancer history **H**= (**P**,**D**). In a Mendelian model, the probability of the person’s genotype given the family cancer history Pr(*G*_0_ | **H**) is the updated population prevalence Pr (*G*_0_) by incorporating family cancer history ***H***. Here, *G*_0_ denotes the genotype of the person of interest, which can be *G*_*m*_, *G*_*f*_, or *G*_*c*_. We can estimate it via the following formula:

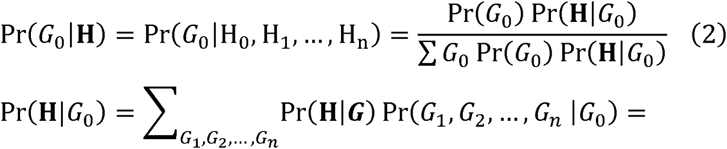

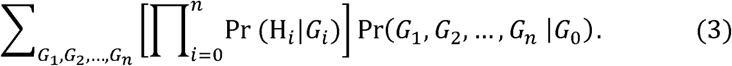

In equations 2 and 3, *n* is the total number of the counselee’s relatives within a family. Pr (**H**|*G*_0_) is the probability of the phenotypes for the whole pedigree given the genotype of the counselee, which is the weighted average of the probabilities of family history given each possible genotype configuration of all relatives Pr (**H**|G). The weights are the probabilities of the genotype configuration based on Mendelian transmission. Pr (**H**|G) are products of the individual probability distributions of penetrance Pr (*H*_*i*_ |*G*_*i*_) when we assume conditional independence. With the equations above for Pr (*G*_0_ |**H**), we can then calculate the four probabilities in equation (1), respectively, by assigning the right person as *G*_0_ and assuming a known genotype status in the parents, when needed.

The posterior probability calculation is performed using the Elston-Stewart peeling algorithm. This algorithm characterizes Mendelian transmission using a transmission matrix of the probability of the genotype for an individual, given the genotypes of the father and mother, Pr (*G*_*i*_ |*G*_*fi*_, *G*_*mi*_) (Fernando et al. 1993).

With the calculated probabilities, we classify the DNM status based on a cutoff on the probability. Pr(*G*_*c*_ is *de novo* | *G*_*c*_ is germline,**D**,**P**) ≥ *cutoff* are identified as DNM carriers. We used the validation data in this study to determine an ideal cutoff.

For Famdenovo.TP53, we used a previous penetrance estimate for *TP53* mutation carriers and noncarriers from six large pediatric sarcoma families at MD Anderson, which are not included in this study (Wu et al. 2006). We assumed the *TP53* mutation follows the Hardy-Weinberg equilibrium, but this could be modified by user input when homozygous genotype information is published. The mutation prevalence, i.e., allele frequency, is specified as 0.0006 for pathogenic *TP53* mutations, which was derived in our previous study (Peng et al. 2017). The assumed frequencies for the three genotypes (homozygous reference, heterozygous, and homozygous variant) were 0.9988, 0.001199, and 3.6×10^−07^, respectively. We used 20% as percent DNMs among all mutations. The allele frequency and DNM rate are priors that are then updated by family history to generate a posterior probability. Both penetrance and prevalence estimates were previously validated using external study cohorts that are different from those in this study (Peng et al. 2017).

For Famdenovo.BRCA, we used previous externally validated penetrance estimates of *BRCA1* and *BRCA2* mutations of the US population for both Ashkenazi Jewish (AJ) and non-AJ families (Parmigiani et al. 2007; Chen et al. 2006a). We used 0.00305 and 0.0034 as the mutation allele frequencies for *BRCA1* and *BRCA*2, respectively. We assumed a 15% DNM rate among all mutations. The allele frequency and DNM rate are priors that were then updated by family history to generate a posterior probability. The same cutoff value of 0.15 was used to classify DNMs in *BRCA1/2*.

### Study cohorts

We evaluated our method on *TP53* using four cohorts (**Table S12**). (A) Families with LFS primarily ascertained through clinical criteria were included in the MD Anderson cohort. Details on data collection, specific cancers, and germline testing for this cohort have been published (Shin et al. 2020). This cohort has 82 families with known *de novo* status and 58 families with unknown *de novo* status. (B) The National Cancer Institute (NCI) LFS cohort (NCT01443468), our second cohort, is from a long-term prospective, natural history study that started in 2011 (Mai et al. 2016). The cohort includes individuals meeting classic or Li-Fraumeni-like diagnostic criteria, having a pathogenic germline *TP53* mutation or a first- or second-degree relative with a *TP53* mutation, or having a personal history of choroid plexus carcinoma, adrenocortical carcinoma, or at least three primary cancers (Birch et al. 1994). This cohort includes 66 families with known *de novo* status and 12 families with unknown *de novo* status. (C) The DFCI LFS cohort, our third cohort, is comprised of patients identified in the course of clinical genetics practice. The early years of the cohort were the result of single gene testing of individuals with features suggestive of LFS. Since 2012, the cohort has included individuals with *TP53* mutations identified on multigene panel testing and individuals referred for clinical consultation of enrollment in a whole-body MRI surveillance protocol. This cohort has 30 families with known *de novo* status and 61 families with unknown *de novo* status. (D) Data from the CHOP were obtained through the Cancer Predisposition Program, which provides care and counseling for children with a genetic predisposition to cancer. This cohort has eight families with known *de novo* status and seven families with unknown *de novo* status.

We also tested Famdenovo.BRCA using a small set of family data that is from the Cancer Genetics Network (CGN) (Anton-Culver et al. 2003), http://www.cancergen.org/studies.shtml, a network established by NCI in 1999 to explore the impact of genetics on cancer susceptibility. Individuals with a family history of cancer were enrolled at clinical centers across the US and completed questionnaires regarding family history of cancer and other relevant information. Most families were ascertained through high-risk clinics and have a family history of breast or ovarian cancers.

### Mutation testing

For the MD Anderson cohort, peripheral blood samples were collected after informed consents were obtained. The probands’ *TP53* mutation status was determined by PCR sequencing of exons 2-11 (Hwang et al. 2003). At MD Anderson Cancer Center, when a *TP53* mutation was identified, all first-degree relatives of the proband (affected and unaffected by cancer) and any other family member at risk of carrying the familial mutation were tested. Extending germline testing based on mutation status and not on phenotype of family members should not introduce ascertainment bias during analysis (Katki et al. 2008). Individuals unavailable for testing (largely deceased) linked to mutation carriers were considered as obligate mutation carriers. No other family member was tested when the proband tested negative.

For the NCI cohort, copies of the clinical *TP53* test reports were obtained and verified by the study team for those tested prior to enrollment. For individuals actively participating in the protocol and not previously tested, clinical genetic testing was performed after enrollment. All at-risk family members of individuals who tested positive for a mutation (either prior to enrollment or on study) were offered the option of having site-specific testing through the study. No testing was offered to relatives if the proband tested negative for a *TP53* mutation. Additionally, high resolution melt analysis and multiplex ligation-dependent probe amplification were performed in the NCI cohort to detect large deletions or genomic rearrangements.

For the DFCI cohort, the patient information was collected by searching through the Clinical Operations and Research Information System (CORIS) database at the DFCI. The classic (Li and Fraumeni Jr. 1969) and updated Chompret (Tinat et al. 2009) criteria were applied for selecting eligible families. The methods for *TP53* testing were further described in a previous study from DFCI (Rath et al. 2013).

For the CHOP cohort, pediatric oncology patients were evaluated in the Cancer Predisposition Program based on a primary tumor type concordant with the LFS tumor spectrum (e.g., adrenocortical carcinoma, choroid plexus carcinoma, early onset rhabdomyosarcoma, multiple primary cancers, etc.) and/or a suggestive family cancer history. Most patients met classic or Chompret criteria. The laboratories used to complete clinical *TP53* testing included the Genetic Diagnostic Laboratory at the University of Pennsylvania, The Hospital for Sick Children, or Ambry Genetics.

### Model evaluation and comparison

In four cohorts (described above) of *TP53* mutation carriers with known *de novo* status, we used Famdenovo.TP53 to calculate the *de novo* probability for each individual. Because mutation carriers within the same family are not independent, we used family-wise *de novo* probability and status for model evaluation. If a family had at least one family member with a *de novo* probability that was over the cut-off, they were defined as a *de novo* family; otherwise they were defined as a familial family.

Validation was performed on 186 families with known DNM status from the four cohorts. We used OEs to evaluate the calibration and ROC curves to evaluate our model’s discrimination ability. The OE is the ratio between the observed number of *de novo TP53* mutations and the summation of the estimated probabilities of *de novo TP53* mutations; ideally, the observed number of DNMs equals the estimated number (OE = 1). The ROC curve is generated by plotting the true positive rate against the false positive rate at various prediction cutoffs using Famdenovo.TP53. A high area under the ROC curve (AUC), i.e., concordance index, indicates that we can find a point on the ROC curve for determining the *de novo* status with a high true positive rate and a low false positive rate.

We also compared our method with the other two criterion-based prediction models. The partial AUC is rescaled to the full range of 0 to 1 using the following equation:

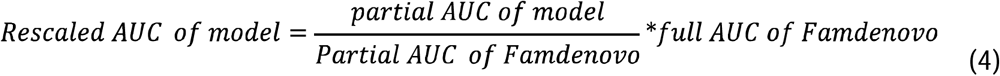

Finally, 95% CI were obtained by bootstrapping the AUC values of Famdenovo and the AUC or rescaled AUC of the two criterion-based models.

DATA ACCESS

### Open Access Data and Script

*TP53* mutation and cancer data from all studies: Table S1.

*BRCA1/2* mutation and cancer data from CGN: Table S11.

R package: Famdenovo_0.1.1.tar.gz.

Supplementary codes: http://bioinformatics.mdanderson.edu/public-software/famdenovo.

### Controlled Access Data

*TP53* family history data can be requested from the LiFE consortium:

MDA: contact Dr. Louise Strong at, Dr. Wenyi Wang at.

NCI: contact Dr. Sharon Savage at. DFCI: contact Dr. Judy Garber at.

CHOP: contact Kristin Zelley at, Dr. Wenyi Wang at.

*BRCA1/2* family history data can be requested from the Cancer Genetics Network: contact Dr. Dianne Finkelstein at.

## ACKNOWLEDGEMENTS

We thank Dr. Chris Amos, Dr. Banu Arun and the reviewers for their constructive comments on this manuscript. We thank Timothy Cheng, Jessica Swann for reading the manuscript and provide word editing. We thank the reviewers for their constructive comments which have substantially improved the paper. The work of X. P. and W. W. are supported in part by NIH R01CA158113. E. D., F. G., and W. W. are supported in part by NIH R01CA183793 and R01CA239342. The work of W. W. is also supported in part by NIH P30CA016672. The work of S. A. S. is supported by the intramural research program of the Division of Cancer Epidemiology and Genetics, National Cancer Institute. J. B. and L. C. S. are supported in part by the NIH P01CA34936. The CGN data collection and curation is supported by N01PC74400-6-0-1.

## AUTHOR CONTRIBUTIONS

Concept and design: F. Gao, X. Pan, W. Wang Development of methodology: F. Gao, X. Pan, W. Wang Acquisition of data: J. Bojadzieva, L. C. Strong, P. L. Mai, S. A. Savage, K. Zelley, K .E. Nichols, J. Garber Analysis and interpretation of data: F. Gao, X. Pan, E. B. Dodd-Eaton, C. Vera Recio, M. D. Montierth, D. Braun, V. E. Johnson, W. Wang Writing, review and/or revision of the manuscript: F. Gao, X. Pan, E. B. Dodd-Eaton, C. Vera Recio, J. Bojadzieva, D. Braun, P. L. Mai, V.E. Johnson, K. Zelley, K. E. Nichols, J. E. Garber, S. A. Savage, L. C. Strong, W. Wang Administrative, technical, or material support: E. B. Dodd-Eaton Study Supervision: W. Wang

## REFERENCES

Acuna-Hidalgo R, Veltman JA, Hoischen A. 2016. New insights into the generation and role of de novo mutations in health and disease. Genome Biol 17: 241. https://doi.org/10.1186/s13059-016-1110-1.

Anton-Culver H, Ziogas A, Bowen D, Finkelstein D, Griffin C, Hanson J, Isaacs C, Kasten-Sportes C, Mineau G, Nadkarni P, et al. 2003. The Cancer Genetics Network: Recruitment results and pilot studies. Community Genet 6: 171–177. https://doi.org/10.1159/000078165.

Battle A, Montgomery SB. 2014. Determining causality and consequence of expression quantitative trait loci. Hum Genet 133: 727–735. https://doi.org/10.1007/s00439-014-1446-0.

Birch JM, Hartley AL, Tricker KJ, Prosser J, Condie A, Kelsey AM, Harris M, Jones PH, Binchy A, Crowther D, et al. 1994. Prevalence and diversity of constitutional mutations in the p53 gene among 21 Li-Fraumeni families. Cancer Res 54: 1298–1304.

Chen S, Iversen ES, Friebel T, Finkelstein D, Weber BL, Eisen A, Peterson LE, Schildkraut JM, Isaacs C, Peshkin BN, et al. 2006a. Characterization of BRCA1 and BRCA2 mutations in a large United States sample. J Clin Oncol.

Chen S, Wang W, Broman KW, Katki HA, Parmigiani G. 2004. BayesMendel: an R environment for Mendelian risk prediction. Stat Appl Genet Mol Biol 3: 1–19. https://doi.org/10.2202/1544-6115.1063.

Chen S, Wang W, Lee S, Nafa K, Lee J, Romans K, Watson P, Gruber SB, Euhus D, Kinzler KW, et al. 2006b. Prediction of germline mutations and cancer risk in the Lynch syndrome. JAMA 296: 1479–1487. https://doi.org/10.1001/jama.296.12.1479.

Conrad DF, Jakobsson M, Coop G, Wen X, Wall JD, Rosenberg NA, Pritchard JK. 2006. A worldwide survey of haplotype variation and linkage disequilibrium in the human genome. Nat Genet 38: 1251–1260. https://doi.org/10.1038/ng1911.

Correa H. 2016. Li-Fraumeni Syndrome. J Pediatr Genet 05(02): 084–088. doi: 10.1055/s-0036-1579759.

De Ligt J, Willemsen MH, Van Bon BWM, Kleefstra T, Yntema HG, Kroes T, Vulto-van Silfhout AT, Koolen DA, De Vries P, Gilissen C, et al. 2012. Diagnostic exome sequencing in persons with severe intellectual disability. N Engl J Med 367: 1921–1929. https://doi.org/10.1056/NEJMoa1206524.

De Rubeis S, He X, Goldberg AP, Poultney CS, Samocha K, Cicek AE, Kou Y, Liu L, Fromer M, Walker S, et al. 2014. Synaptic, transcriptional and chromatin genes disrupted in autism. Nature 515: 209–215. https://doi.org/10.1038/nature13772.

Dryja TP, Morrow JF, Rapaport JM. 1997. Quantification of the paternal allele bias for new germline mutations in the retinoblastoma gene. Hum Genet 100: 446–449. https://doi.org/10.1007/s004390050531.

Euhus DM, Smith KC, Robinson L, Stucky A, Olopade OI, Cummings S, Garber JE, Chittenden A, Mills GB, Rieger P, et al. 2002. Pretest prediction of BRCA1 or BRCA2 mutation by risk counselors and the computer model BRCAPRO. J Natl Cancer Inst 94: 844–851. https://doi.org/10.1093/jnci/94.11.844.

Evans DG, Howard E, Giblin C, Clancy T, Spencer H, Huson SM, Lalloo F. 2010. Birth incidence and prevalence of tumor-prone syndromes: estimates from a UK family genetic register service. Am J Med Genet A 152A: 327–332. https://doi.org/10.1002/ajmg.a.33139.

Fernando RL, Stricker C, Elston RC. 1993. An efficient algorithm to compute the posterior genotypic distribution for every member of a pedigree without loops. Theor Appl Genet Int J Plant Breed Res 87: 89–93. https://doi.org/10.1007/BF00223750.

Francioli LC, Polak PP, Koren A, Menelaou A, Chun S, Renkens I, Van Duijn CM, Swertz M, Wijmenga C, Van Ommen G, et al. 2015. Genome-wide patterns and properties of de novo mutations in humans. Nat Genet 47: 822–826. http://dx.doi.org/10.1038/ng.3292.

Goldmann JM, Veltman JA, Gilissen C. 2019. De Novo Mutations Reflect Development and Aging of the Human Germline. Trends Genet.

Gonzalez KD, Buzin CH, Noltner KA, Gu D, Li W, Malkin D, Sommer SS. 2009. High frequency of de novo mutations in Li-Fraumeni syndrome. J Med Genet 46: 689–693. https://doi.org/10.1136/jmg.2008.058958.

Hwang SJ, Lozano G, Amos CI, Strong LC. 2003. Germline p53 mutations in a cohort with childhood sarcoma: sex differences in cancer risk. Am J Hum Genet 72: 975–983. https://doi.org/10.1086/374567.

Iossifov I, O’Roak BJ, Sanders SJ, Ronemus M, Krumm N, Levy D, Stessman HA, Witherspoon KT, Vives L, Patterson KE, et al. 2014. The contribution of de novo coding mutations to autism spectrum disorder. Nature 515: 216–221. https://doi.org/10.1038/nature13908.

Jett K, Friedman JM. 2010. Clinical and genetic aspects of neurofibromatosis 1. Genet Med 12: 1–11. https://doi.org/10.1097/GIM.0b013e3181bf15e3.

Jónsson H, Sulem P, Arnadottir GA, Pálsson G, Eggertsson HP, Kristmundsdottir S, Zink F, Kehr B, Hjorleifsson KE, Jensson B, et al. 2018. Multiple transmissions of de novo mutations in families. Nat Genet 50: 1674–1680. https://doi.org/10.1038/s41588-018-0259-9.

Katki HA, Blackford A, Chen S, Parmigiani G. 2008. Multiple diseases in carrier probability estimation: accounting for surviving all cancers other than breast and ovary in BRCAPRO. Stat Med 27: 4532–4548. https://doi.org/10.1002/sim.3302.

Kondrashov AS. 2003. Direct estimates of human per nucleotide mutation rates at 20 loci causing Mendelian diseases. Hum Mutat 21: 12–27. https://doi.org/10.1002/humu.10147.

Kong A, Frigge ML, Masson G, Besenbacher S, Sulem P, Magnusson G, Gudjonsson SA, Sigurdsson A, Jonasdottir A, Wong WS, et al. 2012. Rate of de novo mutations and the importance of father’s age to disease risk. Nature 488: 471–475. https://doi.org/10.1038/nature11396.

Krumm N, Turner TN, Baker C, Vives L, Mohajeri K, Witherspoon K, Raja A, Coe BP, Stessman HA, He ZX, et al. 2015. Excess of rare, inherited truncating mutations in autism. Nat Genet 47: 582–588. https://doi.org/10.1038/ng.3303.

Leroy B, Ballinger ML, Baran-Marszak F, Bond GL, Braithwaite A, Concin N, Donehower LA, El-Deiry WS, Fenaux P, Gaidano G, et al. 2017. Recommended guidelines for validation, quality control, and reporting of TP53 variants in clinical practice. Cancer Res 77: 1250–1260. https://doi.org/10.1158/0008-5472.CAN-16-2179.

Li FP, Fraumeni Jr. JF. 1969. Soft-tissue sarcomas, breast cancer, and other neoplasms. A familial syndrome? Ann Intern Med 71: 747–752.

Lipson M, Loh PR, Sankararaman S, Patterson N, Berger B, Reich D. 2015. Calibrating the Human Mutation Rate via Ancestral Recombination Density in Diploid Genomes. PLoS Genet 11: e1005550. https://doi.org/10.1371/journal.pgen.1005550.

Lynch M. 2010. Rate, molecular spectrum, and consequences of human mutation. Proc Natl Acad Sci U S A 107: 961–968. https://doi.org/10.1073/pnas.0912629107.

Mai PL, Best AF, Peters JA, DeCastro RM, Khincha PP, Loud JT, Bremer RC, Rosenberg PS, Savage SA. 2016. Risks of first and subsequent cancers among TP53 mutation carriers in the National Cancer Institute Li-Fraumeni syndrome cohort. Cancer 122: 3673–3681. https://doi.org/10.1002/cncr.30248.

Malekzadeh P, Rosenberg SA, Drew C, Invest JC, Malekzadeh P, Pasetto A, Robbins PF, Parkhurst MR, Paria BC, Jia L, et al. 2019. Neoantigen screening identifies broad TP53 mutant immunogenicity in patients with epithelial cancers. J Clin Invest 129. https://doi.org/10.1172/JCI123791.

Malkin D, Li FP, Strong LC, Fraumeni Jr. JF, Nelson CE, Kim DH, Kassel J, Gryka MA, Bischoff FZ, Tainsky MA, et al. 1990. Germ line p53 mutations in a familial syndrome of breast cancer, sarcomas, and other neoplasms. Science (80-) 250: 1233–1238.

Michaelson JJ, Shi Y, Gujral M, Zheng H, Malhotra D, Jin X, Jian M, Liu G, Greer D, Bhandari A, et al. 2012. Whole-genome sequencing in autism identifies hot spots for de novo germline mutation. Cell 151: 1431–1442. https://doi.org/10.1016/j.cell.2012.11.019.

Nachman MW, Crowell SL. 2000. Estimate of the mutation rate per nucleotide in humans. Genetics 156: 297–304.

Nichols KE, Malkin D, Garber JE, Fraumeni J, Li FP. 2001. Germ-line p53 mutations predispose to a wide spectrum of early-onset cancers. Cancer Epidemiol Biomarkers Prev 10: 83–87.

Olivier M, Goldgar DE, Sodha N, Ohgaki H, Kleihues P, Hainaut P, Eeles RA. 2003. Li-Fraumeni and Related Syndromes. Cancer Res 63: 6643–6650.

Parmigiani G, Berry D, Aguilar O. 1998. Determining carrier probabilities for breast cancer-susceptibility genes BRCA1 and BRCA2. Am J Hum Genet 62: 145–158. https://doi.org/10.1086/301670.

Parmigiani G, Chen S, Iversen Jr. ES, Friebel TM, Finkelstein DM, Anton-Culver H, Ziogas A, Weber BL, Eisen A, Malone KE, et al. 2007. Validity of models for predicting BRCA1 and BRCA2 mutations. Ann Intern Med 147: 441–450. https://doi.org/10.7326/0003-4819-147-7-200710020-00002.

Peng G, Bojadzieva J, Ballinger ML, Li J, Blackford AL, Mai PL, Savage SA, Thomas DM, Strong LC, Wang W. 2017. Estimating TP53 mutation carrier probability in families with li-fraumeni syndrome using LFSPRO. Cancer Epidemiol Biomarkers Prev 26: 837–844. https://doi.org/10.1158/1055-9965.EPI-16-0695.

Rahbari R, Wuster A, Lindsay SJ, Hardwick RJ, Alexandrov LB, Al Turki S, Dominiczak A, Morris A, Porteous D, Smith B, et al. 2016. Timing, rates and spectra of human germline mutation. Nat Genet 48: 126–133. https://doi.org/10.1038/ng.3469.

Rath MG, Masciari S, Gelman R, Miron A, Miron P, Foley K, Richardson AL, Krop IE, Verselis SJ, Dillon DA, et al. 2013. Prevalence of germline TP53 mutations in HER2+ breast cancer patients. Breast Cancer Res Treat 139: 193–198. https://doi.org/10.1007/s10549-012-2375-z.

Renaux-Petel M, Charbonnier F, Thery JC, Fermey P, Lienard G, Bou J, Coutant S, Vezain M, Kasper E, Fourneaux S, et al. 2018. Contribution of de novo and mosaic TP53 mutations to Li-Fraumeni syndrome. J Med Genet 55: 173–180. https://doi.org/10.1136/jmedgenet-2017-104976.

Robinson EB, Samocha KE, Kosmicki JA, McGrath L, Neale BM, Perlis RH, Daly MJ. 2014. Autism spectrum disorder severity reflects the average contribution of de novo and familial influences. Proc Natl Acad Sci U S A 111: 15161–15165. https://doi.org/10.1073/pnas.1409204111.

Shin SJ, Dodd-Eaton EB, Gao F, Bojadzieva J, Chen J, Kong X, Amos CI, Ning J, Strong LC, Wang W. 2020. Penetrance estimates over time to first and second primary cancer diagnosis in families with Li-Fraumeni syndrome: A single institution perspective. Cancer Res 80(2):347–353.

Tikkanen T, Leroy B, Fournier JL, Risques RA, Malcikova J, Soussi T. 2018. Seshat: A Web service for accurate annotation, validation, and analysis of TP53 variants generated by conventional and next-generation sequencing. Hum Mutat 39: 925–933. https://doi.org/10.1002/humu.23543.

Tinat J, Bougeard G, Baert-Desurmont S, Vasseur S, Martin C, Bouvignies E, Caron O, Paillerets BB, Berthet P, Dugast C, et al. 2009. 2009 Version of the Chompret Criteria for Li Fraumeni Syndrome. J Clin Oncol 27: e108–e109. https://doi.org/10.1200/jco.2009.22.7967.

Walerych D, Napoli M, Collavin L, Del Sal G. 2012. The rebel angel: Mutant p53 as the driving oncogene in breast cancer. Carcinogenesis 33: 2007–2017.

Wang W, Chen S, Brune KA, Hruban RH, Parmigiani G, Klein AP. 2007. PancPRO: risk assessment for individuals with a family history of pancreatic cancer. J Clin Oncol 25: 1417–1422. https://doi.org/10.1200/JCO.2006.09.2452.

Wang W, Niendorf KB, Patel D, Blackford A, Marroni F, Sober AJ, Parmigiani G, Tsao H. 2010. Estimating CDKN2A carrier probability and personalizing cancer risk assessments in hereditary melanoma using MelaPRO. Cancer Res 70: 552–559. https://doi.org/10.1158/0008-5472.CAN-09-2653.

Weitzel JN, Chao EC, Nehoray B, Van Tongeren LR, LaDuca H, Blazer KR, Slavin T, Facmg DABMD, Pesaran T, Rybak C, et al. 2018. Somatic TP53 variants frequently confound germ-line testing results. Genet Med 20: 809–816. https://doi.org/10.1038/gim.2017.196.

Wong WSW, Solomon BD, Bodian DL, Kothiyal P, Eley G, Huddleston KC, Baker R, Thach DC, Iyer RK, Vockley JG, et al. 2016. New observations on maternal age effect on germline de novo mutations. Nat Commun 7: 1–10. http://dx.doi.org/10.1038/ncomms10486.

Wu CC, Shete S, Amos CI, Strong LC. 2006. Joint effects of germ-line p53 mutation and sex on cancer risk in Li-Fraumeni syndrome. Cancer Res 66: 8287–8292.

