## Supplementary Figure 1 for "A pedigree-based prediction model identifies carriers of deleterious *de novo* mutations in families with Li-Fraumeni syndrome"

**Supplementary Figure 1.** Distributions of pedigree sizes in the LFS dataset and the CGN-BRCA dataset.

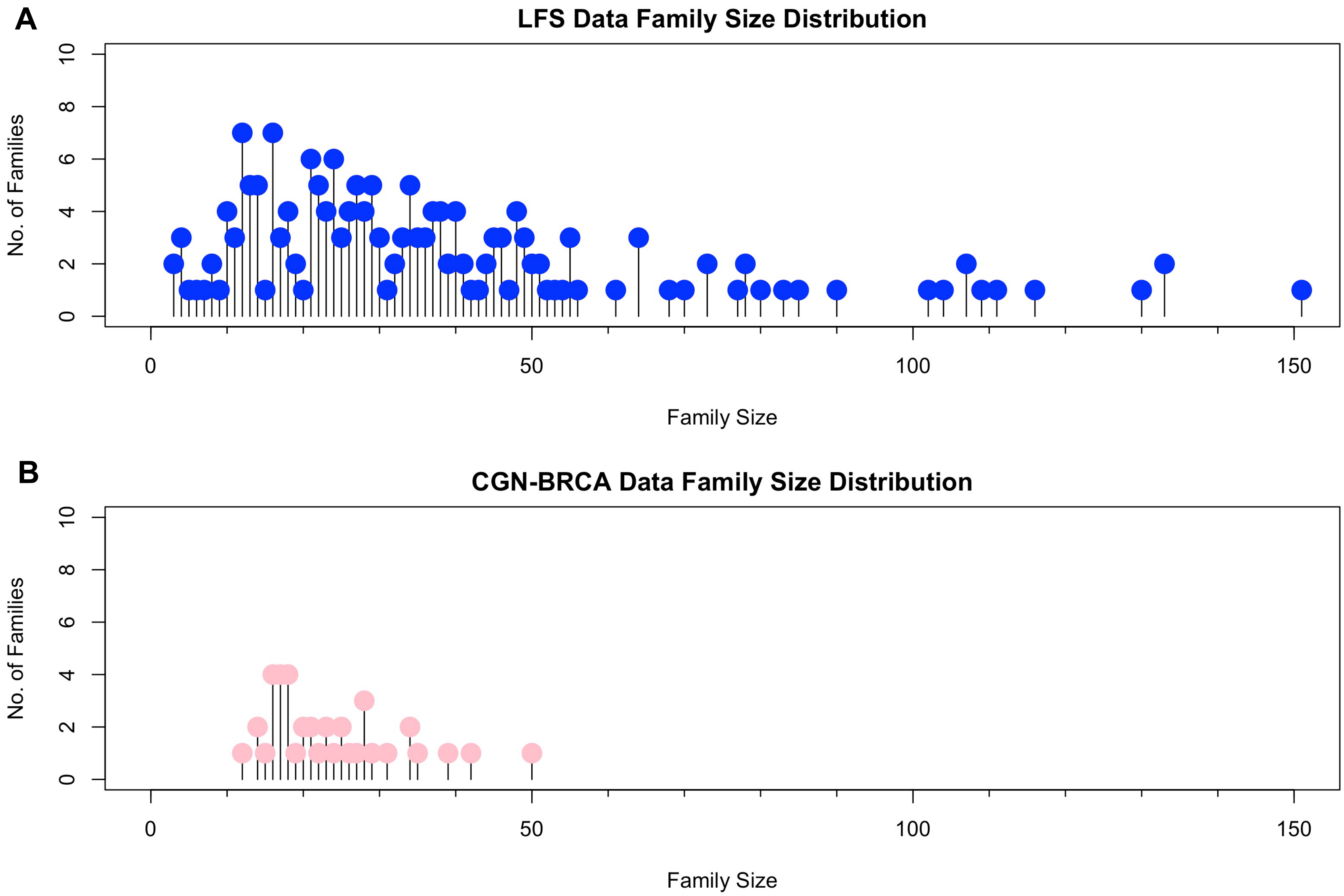
