## Supplementary Figure 2 for "A pedigree-based prediction model identifies carriers of deleterious *de novo* mutations in families with Li-Fraumeni syndrome"

**Supplementary Figure 2.** Venn diagrams of the mutations observed in the validation and discovery sets. The blue circles enclose the mutations in the validation set and the orange circles enclose the mutations in the discovery set. The amino acid change caused by the *de novo* mutations that occurred in both validation and discovery set is shown inside brackets. The numbers inside parenthesis are the frequency of the mutation in the validation and discovery set, respectively. The mutations in red font in Venn diagram (**B**) cause the amino acid changes p.R248Q and p.R248W in the *p53* protein.

**A.** *de novo* mutations in the validation and discovery set

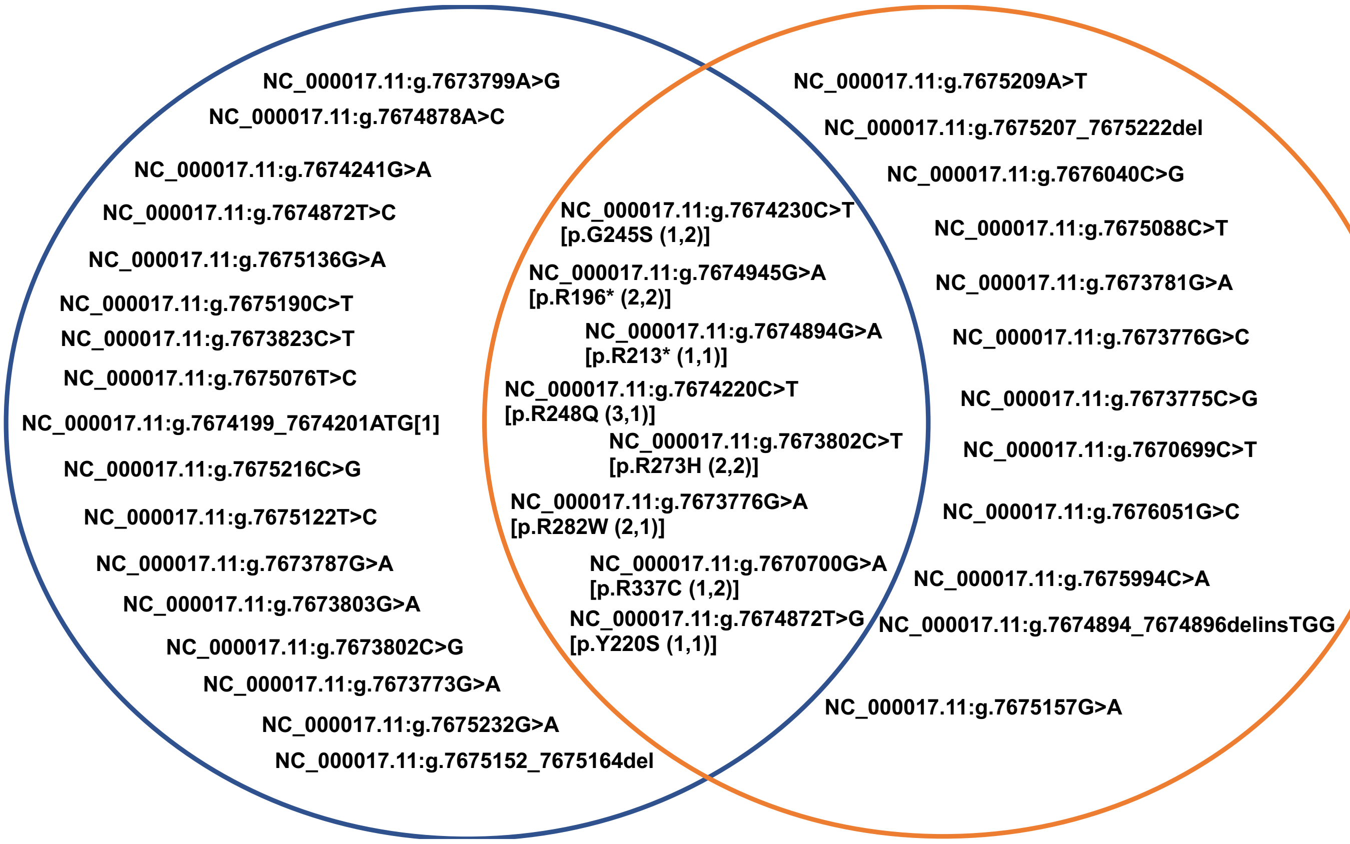


**B.** Familial mutations in the validation and discovery set

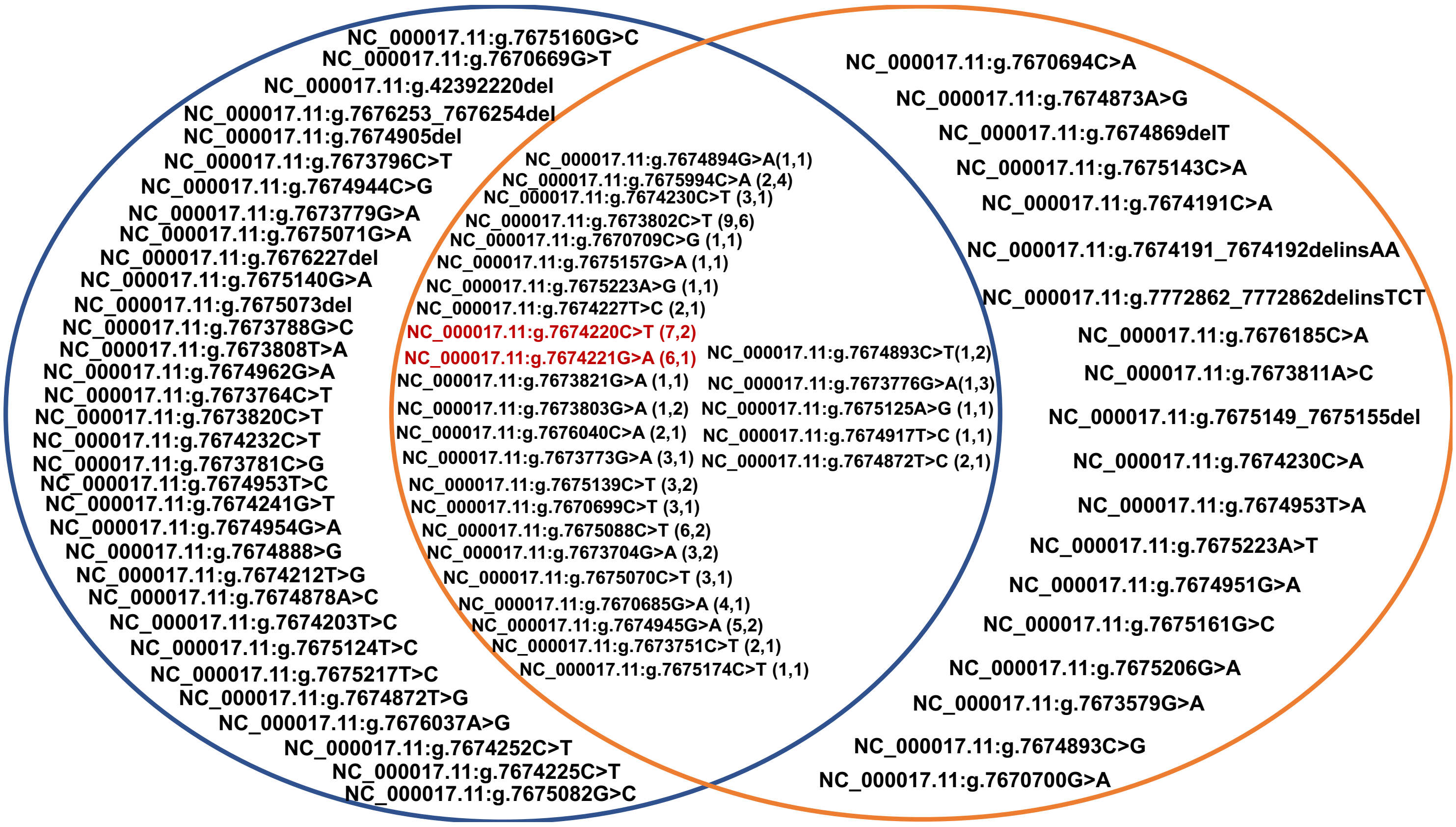
