## Supplementary Figure 3 for "A pedigree-based prediction model identifies carriers of deleterious *de novo* mutations in families with Li-Fraumeni syndrome"

**Supplementary Figure 3.** Distribution of probabilities of *de novo BRCA1/2* mutations for 66 mutation carriers from the CGN dataset. The gray line denotes a suggested cutoff of 0.15 for the status of *de novo* mutations.

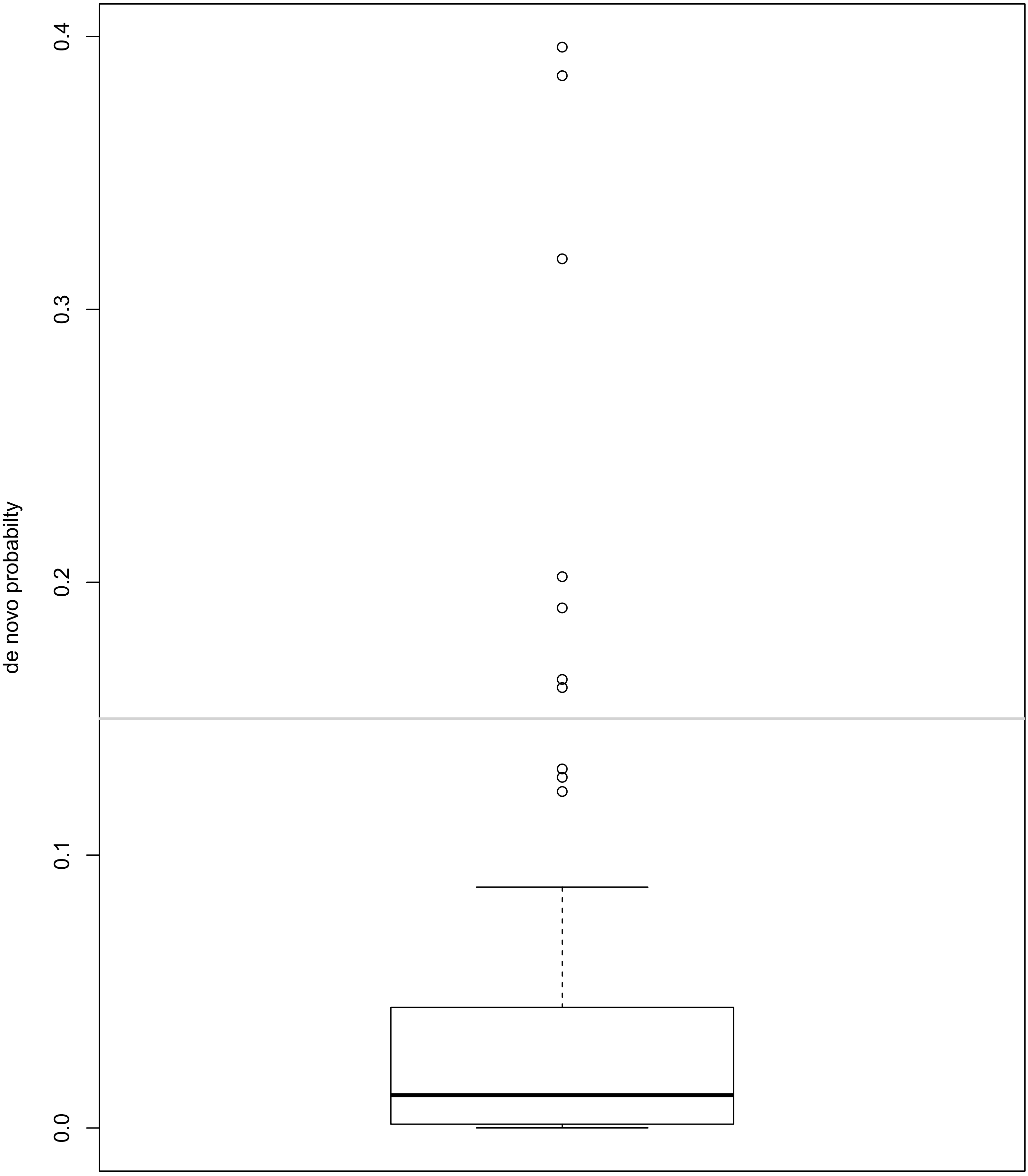
