## Supplementary Figure 4 for "A pedigree-based prediction model identifies carriers of deleterious *de novo* mutations in families with Li-Fraumeni syndrome"

**Supplementary Figure 4.** Pedigree plots for the 7 predicted DNMs in *BRCA1/2*. The blue square highlights the family member predicted to be DNM carrier. The cancer type and age at diagnosis (red font) are shown under each cancer patient (black filled circle or square). The age at last contact is also listed whenever available for each family member.

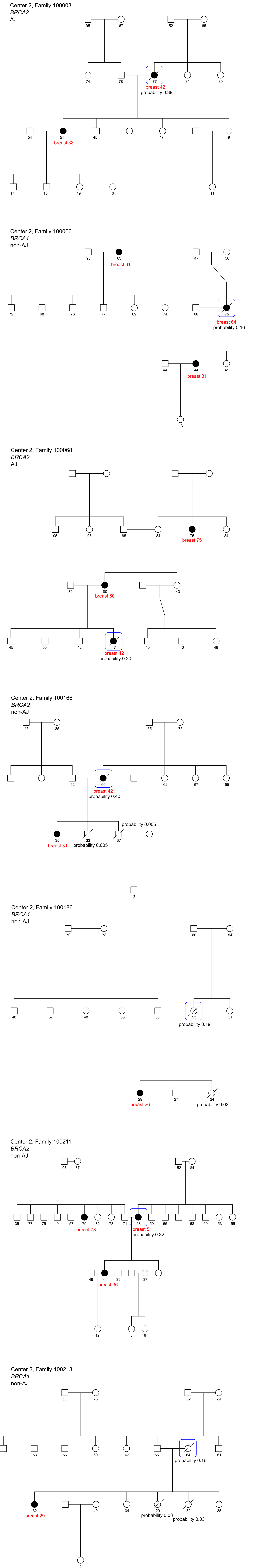
