## Supplementary Table 2 for "A pedigree-based prediction model identifies carriers of deleterious *de novo* mutations in families with Li-Fraumeni syndrome"

**Supplementary Table 2.** Sensitivity and specificity of Famdenovo under different cutoffs for classifying DNM status of *TP53* mutations. The dataset used for evaluating sensitivity and specificity is the same as in **Figure 1**, which contains 186 families. There are 42 of these families that have an individual who carries a *de novo* *TP53* mutation. The first column is the cutoff chosen to classify the DNM status as 1 if the probability is greater than the cutoff, and 0 otherwise. The second column is the number of predicted positives using the corresponding cutoff. The third column is the number of true positives out of those that are predicted. The fourth and fifth columns are the corresponding sensitivity and specificity, respectively.

| **Prediction cutoff** | **Called positives, no.** | **Called true positives, no.** | **Sensitivity** | **Specificity** |
| --- | --- | --- | --- | --- |
| 0.3 | 37 | 30 | 0.71 | 0.95 |
| 0.2 | 42 | 32 | 0.76 | 0.93 |
| 0.1 | 45 | 33 | 0.79 | 0.92 |
| 0.05 | 55 | 37 | 0.88 | 0.88 |
