## Supplementary Table 3 for "A pedigree-based prediction model identifies carriers of deleterious *de novo* mutations in families with Li-Fraumeni syndrome"

**Supplementary Table 3.** AUC and OE of Famdenovo under different parameter settings. The data used here is the same as in **Figure 1**, which contains 186 families. The MAF parameter refers to minor allele frequency, which is the frequency of mutated alleles in *TP53*. The mRate parameter denotes the expected *de novo* mutation rate in *TP53*. Both parameters correspond to prior values that will be updated in Mendelian models by family history. The row in grey corresponds to the default parameter settings, as recommended by LFSPRO for *TP53* mutations.

| MAF | mRate | AUC (95% CI) | OE (95% CI) |
| --- | --- | --- | --- |
| 3e-5 | 6e-6 | 0.952 ([0.920, 0.977]) | 1.343 ([1.102, 1.646]) |
| 1e-4 | 2e-5 | 0.951 ([0.919, 0.997]) | 1.337 ([1.069, 1.641]) |
| 3e-4 | 6e-5 | 0.950 ([0.917, 0.976]) | 1.332 ([1.093, 1.633]) |
| 1e-3 | 2e-4 | 0.947 ([0.914, 0.975]) | 1.324 ([1.084, 1.619]) |
