## Supplementary Table 4 for "A pedigree-based prediction model identifies carriers of deleterious *de novo* mutations in families with Li-Fraumeni syndrome"

**Supplementary Table 4.** Results for the discovery set of 138 families with *TP53* mutations from the four cohorts, with cutoffs of 0.05, 0.1, and 0.3, respectively.

| **Cohort** | **Cutoff = 0.05,**  **predicted *de novo* / total, no. (%)** | **Cutoff = 0.1,**  **predicted *de novo* / total, no. (%)** | **Cutoff = 0.3,**  **predicted *de novo* / total, no. (%)** |
| --- | --- | --- | --- |
| CHOP | 5/7 (71%) | 4/7(57%) | 2/7(29%) |
| DFCI | 10/61 (16%) | 9/61 (15%) | 7/61 (11%) |
| MDA | 27/58 (47%) | 25/58 (43%) | 23/58 (40%) |
| NCI | 9/12 (75%) | 8/12 (67%) | 6/12 (50%) |
| All cohorts | 51/138 (37%) | 46/138 (33%) | 38/138 (28%) |
