## Supplementary Table 5 for "A pedigree-based prediction model identifies carriers of deleterious *de novo* mutations in families with Li-Fraumeni syndrome"

**Supplementary Table 5.** Contingency table of DNM status (*de novo* or familial) and sex in the combined validation and discovery set. (**A**) Contingency table includes 397 cancer patients (people with cancer information, i.e., healthy people excluded). (**B**) Contingency table includes 233 people who are probands with cancer information.

(A)

|  | **Female** | **Male** |
| --- | --- | --- |
| *de novo* | 61 | 19 |
| familial | 204 | 113 |
| odds ratio | 1.78, 95% CI [1.01, 3.13] | |
| p-value | 0.059 | |

(B)

|  | **female** | **male** |
| --- | --- | --- |
| *de novo* | 59 | 19 |
| familial | 115 | 40 |
| odds ratio | 1.08, 95% CI [0.58, 2.03] | |
| p-value | 0.936 | |
