## Supplementary Table 6 for "A pedigree-based prediction model identifies carriers of deleterious *de novo* mutations in families with Li-Fraumeni syndrome"

**Supplementary Table 6.** Contingency table of DNM status (*de novo* or familial) and whether the patient is a breast cancer patient in the combined validation and discovery set. (**A**) Within 233 probands with cancer information. (**B**) Divide the 233 probands with cancer information into three age categories: [0, 20), [20, 35) and 35+. (**C**) Divide the 233 probands with cancer information in the validation set in three age categories. (**D**) Observations were made for probands with breast cancer as the age range changed.

(A)

|  | **breast** | **other** |
| --- | --- | --- |
| *de novo* | 50 | 28 |
| familial | 75 | 80 |
| odds ratio | 1.9, 95% CI [1.09, 3.33] | |
| p-value | 0.033 | |

(B)

|  | **breast**  **[0-20)** | **other**  **[0-20)** | **breast**  **[20,35)** | **other**  **[20,35)** | **breast**  **[35+)** | **other**  **[35+)** |
| --- | --- | --- | --- | --- | --- | --- |
| *de novo* | 0 | 15 | 23 | 4 | 27 | 9 |
| familial | 0 | 32 | 25 | 26 | 50 | 22 |
| [0-20) | NA | | | | | |
| [20,35) | p-value = 0.0029, OR = 5.98, 95% CI [1.81, 19.76] | | | | | |
| [35+) | p-value = 0.66, OR = 1.32, 95% CI [0.53, 3.27] | | | | | |

(C)

|  | **breast**  **[0-20)** | **other**  **[0-20)** | **breast**  **[20,35)** | **other**  **[20,35)** | **breast**  **[35+)** | **other**  **[35+)** |
| --- | --- | --- | --- | --- | --- | --- |
| *de novo* | 0 | 10 | 11 | 3 | 13 | 2 |
| familial | 0 | 25 | 8 | 11 | 28 | 11 |
| [0-20) | NA | | | | | |
| [20,35) | p-value = 0.073, OR = 5.04, 95% CI [1.05, 24.19] | | | | | |
| [35+) | p-value = 0.31, OR = 2.55, 95% CI [0.49, 13.22] | | | | | |

(D)

| **Age range** | **DNM vs FM** | **ratio of DNM** |
| --- | --- | --- |
| 20-30 | 13 vs 15 | 46% |
| 20-40 | 39 vs. 41 | 49% |
| 20-45 | 43 vs. 53 | 45% |
