## Supplementary Table 7 for "A pedigree-based prediction model identifies carriers of deleterious *de novo* mutations in families with Li-Fraumeni syndrome"

**Supplementary Table 7**. Description of the classic LFS and Chompret (2015) criteria.

| **Criteria** | **Description** |
| --- | --- |
| Classic | - Proband: sarcoma age < 45   AND   - First degree relative: any cancer age < 45   AND   - Other first or second degree relative: any cancer age < 45 or a sarcoma at any age |
| Chompret | - Proband diagnosed with - LFS spectrum cancer age < 46 and - First or second degree relative with LFS spectrum cancer (except breast cancer, if the proband has breast cancer) age < 56 or with multiple tumors   OR   - Proband diagnosed with - Multiple primary cancer (except multiple breast cancer), two of which belong to LFS tumor spectrum and the first cancer < 46 year or - Adrenocortical carcinoma, choroid plexus cancer, or rhabdomyosarcoma of embryonal anaplastic subtype, irrespective of family history - Breast cancer before age 31 |
