## Supplementary Table 8 for "A pedigree-based prediction model identifies carriers of deleterious *de novo* mutations in families with Li-Fraumeni syndrome"

**Supplementary Table 8.** Contingency table of DNM status (*de novo* or familial) and multiple primary cancer (MPC), single primary cancer (SPC) in the combined validation and discovery set. (**A**) Within 233 probands with cancer information. (**B**) Divide the 233 probands with cancer information in three age categories: [0, 20), [20, 35) and 35+. (**C**) Divide the probands in the validation set with cancer information in three age categories.

(A)

|  | **MPC** | **SPC** |
| --- | --- | --- |
| *de novo* | 51 | 27 |
| familial | 80 | 75 |
| odds ratio | 1.77, 95% CI [1.01, 3.11] | |
| p-value | 0.0629 | |

(B)

|  | **MPC**  **[0-20)** | **SPC**  **[0-20)** | **MPC**  **[20,35)** | **SPC**  **[20,35)** | **MPC**  **[35+)** | **SPC**  **[35+)** |
| --- | --- | --- | --- | --- | --- | --- |
| *de novo* | 5 | 10 | 17 | 10 | 29 | 7 |
| familial | 10 | 22 | 22 | 29 | 48 | 24 |
| [0-20) | p-value = 1; OR = 1.1; 95% CI [0.3, 4.07] | | | | | |
| [20,35) | p-value = 0.153, OR = 2.24, 95% CI [0.86, 5.84] | | | | | |
| [35+) | p-value = 0.177, OR = 2.07, 95% CI [0.79, 5.41] | | | | | |

(C)

|  | **MPC**  **[0-20)** | **SPC**  **[0-20)** | **MPC**  **[20,35)** | **SPC**  **[20,35)** | **MPC**  **[35+)** | **SPC**  **[35+)** |
| --- | --- | --- | --- | --- | --- | --- |
| *de novo* | 2 | 8 | 10 | 4 | 13 | 2 |
| familial | 7 | 18 | 8 | 11 | 25 | 14 |
| [0-20) | p-value = 1; OR = 0.65; 95% CI [-1.13, 2.44] | | | | | |
| [20,35) | p-value = 0.16, OR = 3.31, 95% CI [0.65, 20.1] | | | | | |
| [35+) | p-value = 0.18, OR = 3.56, 95% CI [0.65, 37.1] | | | | | |
