## Supplementary Table 12 for "A pedigree-based prediction model identifies carriers of deleterious *de novo* mutations in families with Li-Fraumeni syndrome"

**Supplementary Table 12.** Ascertainment criteria of the four cohorts in our study.

| **Cohort** | **Ascertainment criteria** |
| --- | --- |
| CHOP | Pediatric oncology patients were evaluated in the Cancer Predisposition Program based on a primary tumor type concordant with the LFS tumor spectrum (i.e., adrenocortical carcinoma, choroid plexus carcinoma, early onset RMS, multiple primary cancers, etc.) and/or a suggestive family history. |
| DFCI | Comprised of patients identified in the course of clinical genetics practice. The early years of the cohort were the result of single gene testing of individuals with features suggestive of LFS. Since 2012, the cohort has included individuals with *TP53* mutations identified on multigene panel testing, and individuals referred for clinical consultation of enrollment in a whole-body MRI surveillance protocol. |
| MDA | Prospective LFS families primarily ascertained through clinical criteria were used as part of the MD Anderson cohort. |
| NCI | A prospective cohort. Classic LFS or modified criteria, having a pathogenic germline TP53 mutation or a first- or second-degree relative with a mutation, or a personal history of choroid plexus carcinoma, adrenocortical carcinoma, or at least three primary cancers. |
